# NetMix: A network-structured mixture model for reduced-bias estimation of altered subnetworks

**DOI:** 10.1101/2020.01.18.911438

**Authors:** Matthew A. Reyna, Uthsav Chitra, Rebecca Elyanow, Benjamin J. Raphael

## Abstract

A classic problem in computational biology is the identification of *altered subnetworks:* subnetworks of an interaction network that contain genes/proteins that are differentially expressed, highly mutated, or otherwise aberrant compared to other genes/proteins. Numerous methods have been developed to solve this problem under various assumptions, but the statistical properties of these methods are often unknown. For example, some widely-used methods are reported to output very large subnetworks that are difficult to interpret biologically. In this work, we formulate the identification of altered subnetworks as the problem of estimating the parameters of a class of probability distributions which we call the Altered Subset Distribution (ASD). We derive a connection between a popular method, jActiveModules, and the maximum likelihood estimator (MLE) of the ASD. We show that the MLE is *statistically biased*, explaining the large subnetworks output by jActiveModules. We introduce NetMix, an algorithm that uses Gaussian mixture models to obtain less biased estimates of the parameters of the ASD. We demonstrate that NetMix outperforms existing methods in identifying altered subnetworks on both simulated and real data, including the identification of differentially expressed genes from both microarray and RNA-seq experiments and the identification of cancer driver genes in somatic mutation data.

**Availability:** NetMix is available online at https://github.com/raphael-group/netmix.

## 1 Introduction

A standard paradigm in computational biology is to use interaction networks as prior knowledge in the analysis of high-throughput’ omics data, with applications in protein function prediction [79, 73, 65, 25, 18], gene expression [32, 91, 16, 48, 27], germline variants [55, 12, 56, 43, 45], somatic variants in cancer [66, 87, 57, 84, 64, 42], and other data [39, 10, 20, 89, 35, 77, 13, 60]. One classic approach is to identify *active*, or *altered*, subnetworks of an interaction network that contain outlier measurements. The altered subnetwork problem takes as input: (1) an interaction network whose nodes are biological entities (e.g., genes or proteins) and whose edges represent biological interactions (e.g., physical or genetic interactions, co-expression, etc.); and (2) a measurement or score for each node. The goal is to find high-scoring subnetworks that correspond to functionally related or correlated alterations. This problem was introduced in [48] for gene expression analysis, where gene scores were derived from *p*-values of differential expression. [48] developed the jActiveModules algorithm to solve this problem and identify altered subnetworks of differentially expressed genes. Subsequently, [27] introduced heinz as “the first approach that really tackles and solves the original problem raised by [48] to optimality.” jActiveModules and heinz have become widely-used tools with diverse applications; a few recent examples include mass-spectrometry proteomics [51, 58], damaging *de novo* mutations in schizophrenia and other neurological disorders [36, 17], and single-cell RNA-seq [37, 85, 52].

In the past two decades, many algorithms have been developed to identify altered subnetworks in biological data (reviewed in [26, 20, 63, 64]). Each publication describing a new algorithm demonstrates the performance of their algorithm on specific biological datasets, and many of these publications also benchmark their algorithm against existing algorithms on real and/or simulated data. However, few of these publications prove theoretical guarantees for their algorithm’s performance on a well-defined generative model of the data. Thus, the true performance of these algorithms is often unknown. Indeed, recent benchmarking studies (e.g., [40, 9]) of several widely used network algorithms – including jActiveModules and heinz – show considerate disagreement between subnetworks identified by different methods on the same biological datasets. Moreover, these benchmarking studies (and many others) do not compare network algorithms against single-gene tests that do not use the network; thus, the tacit assumption that interaction networks always improve gene prioritization is often not tested.

Separately, many publications in the statistics and machine learning literature investigate the problem of *detecting* whether or not a network contains an anomalous subnetwork, or a *network anomaly*, e.g., [6, 4, 1, 3, 83, 82, 81, 80, 5]. These papers describe specific generative models of network anomalies and use a rigorous hypothesis-testing framework to prove asymptotic results regarding the conditions under which it is possible to detect a network anomaly. Importantly, these papers also provide theoretical guarantees about conditions under which a network contributes to anomaly detection. However, the network anomaly literature does not specifically address the altered subnetwork problem studied in computational biology, with three key differences. First, the *detection* problem of deciding whether or not an altered subnetwork exists is not the same as the *estimation* problem of identifying the nodes in an altered subnetwork. Second, biological networks have a finite size, and it is unclear what guarantees the asymptotic results provide for finite-size networks. Finally, the topological constraints on network anomalies are different from those considered in computational biology.

In this paper, we aim to bridge the gap between the theoretical guarantees in the network anomaly literature and the practical problem of identifying altered subnetworks in biological data. We provide a rigorous formulation of the *Altered Subnetwork Problem*, the problem that jActiveModules [48], heinz [27], and other methods aim to solve. Our formulation of the Altered Subnetwork Problem is inspired by the generative model used in the network anomaly literature, but requires that the altered subnetwork is a connected subnetwork, a constraint motivated by the topology of signaling pathways [11, 50] and by the seminal works of [48] and [27].

We show that the Altered Subnetwork Problem is equivalent to estimating the parameters of a distribution which we define as the *Altered Subset Distribution (ASD)*. We prove that the jActiveModules problem [48] is equivalent to finding a maximum likelihood estimator (MLE) of the parameters of the ASD for connected subgraphs. At the same time, we demonstrate that if (1) the size of the altered subnetwork is moderately small and (2) the scores of nodes inside and outside of the altered subnetwork are not well-separated, then the MLE is a *biased* estimator of the size of the altered subnetwork. This statistical bias provides a rigorous explanation for the large subnetworks produced by jActiveModules [48]. We also show that the size of the altered subnetworks identified by heinz [27] are biased for most choices of its user-defined parameter.

We introduce a new algorithm, NetMix, that combines a Gaussian mixture model and a combinatorial optimization algorithm to identify altered subnetworks. We show that NetMix is a reduced-bias estimator of the size of the altered subnetwork. We demonstrate that NetMix outperforms other methods for identifying altered subnetworks on simulated data, gene expression data, and somatic mutation data.

## 2 Altered Subnetworks, Altered Subsets, and Maximum Likelihood Estimation

### 2.1 Altered Subnetwork Problem

Let *G* = (*V*, *E*) be a biological interaction network with a measurement, or score, *X_v_* for each vertex *v* ∈ *V*. We assume that there is a connected subnetwork *A* in *G*, the *altered subnetwork*, whose scores are derived from a different distribution than the scores of the vertices not in *A* (Figure 1). The goal of the Altered Subnetwork Problem is to find *A*. The problem is defined formally as follows.

**Figure 1:**
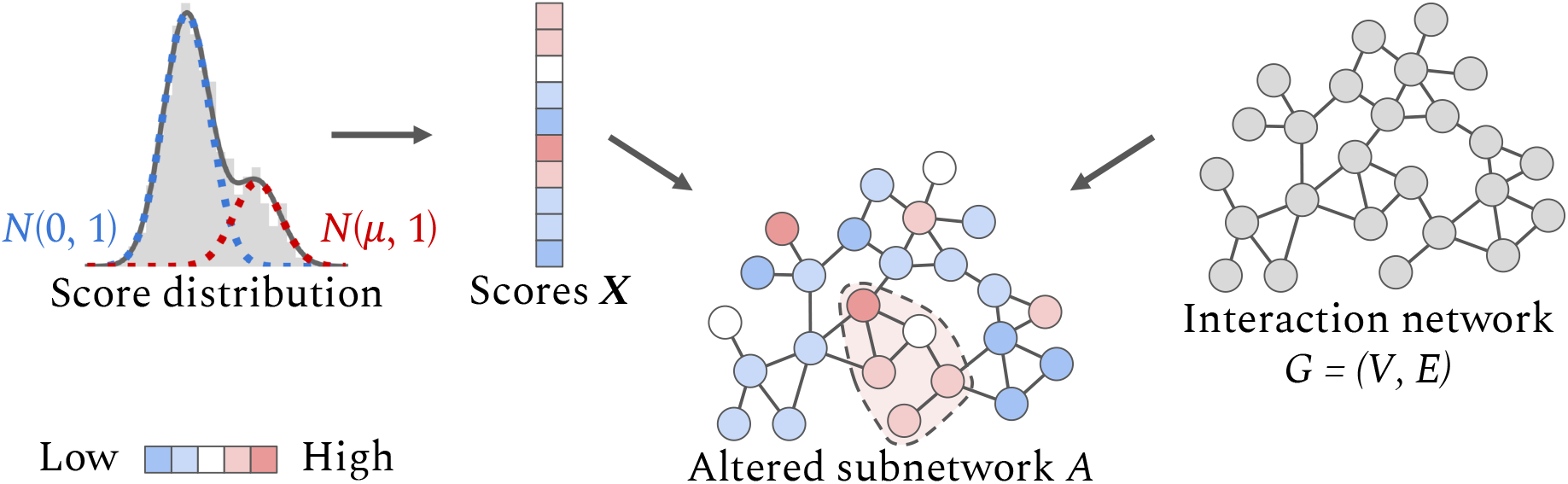
Altered Subnetwork Problem. Measurements, or scores, **X** from a high-throughput experiment are drawn from one of two distributions: genes/proteins in an altered subnetwork *A* of an interaction network *G* = (*V, E*) have scores drawn from an altered distribution *N*(*μ*, 1) with *μ* > 0, while genes/proteins not in *A* have scores drawn from a background distribution *N*(0,1). The difficulty in identifying *A* depends on the separation *μ* between the distributions and the size |*A*| of the altered subnetwork.

#### Altered Subnetwork Problem (ASP)

*Let G* = (*V*, *E*) *be a graph with vertex scores* **X** = (*X_v_*)_*v*∈*V*_, *and let A* ⊆ *V be a connected subgraph of G*. *Suppose that*

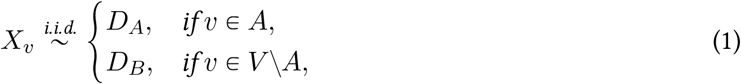

*where D_A_ is the* altered *distribution and D_B_ is the* background *distribution*. *Given G and* **X**, *find A*.

Note that the ASP assumes that the network *G* has a *single* altered subnetwork *A*. When the network has multiple altered subnetworks, one can recursively solve the ASP to identify more than one altered subnetwork.

The seminal algorithm for solving the ASP is jActiveModules [48]. jActiveModules takes as input a *p*-value *p_v_* for each vertex *v*; e.g., a *p*-value of differential gene expression. Under the null hypothesis, the *p*-values *p_v_* across genes are distributed according to the uniform distribution *U*(0,1). jActiveModules transforms the *p*-values into scores *X_v_* = Φ^−1^(1 − *p_v_*), where Φ is the CDF of a standard normal distribution. Thus, jActiveModules solves the ASP with background distribution *D_B_* = *N*(0,1). jActiveModules aims to find a connected subgraph *Â* that maximizes^1^ 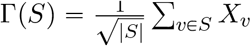, i.e.,

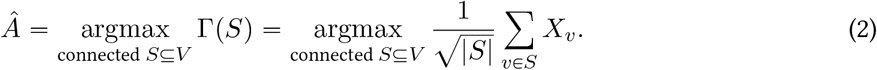

The presentation of jActiveModules in [48] does not specify the altered distribution *D_A_*. However, in Section 2.2, we argue that the choice of the objective function in (2) implicitly assumes that *D_A_* = *N*(*μ*, 1) for some parameter *μ* > 0. Thus, we define the normally distributed ASP as follows.

#### Normally Distributed Altered Subnetwork Problem

*Let G* = (*V*, *E*) *be a graph with vertex scores* **X** = (*X_v_*)_*v*∈*V*_, *and let A* ⊆ *V be a connected subgraph of G*. *Suppose that for some μ* > 0,

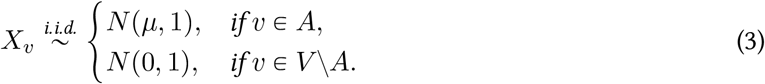

*Given G and* **X**, *find A*.

The Normally Distributed ASP has a sound statistical interpretation: if the *p*-values *p_v_* of the genes are derived from an asymptotically normal test statistic, as is often the case, then the transformed *p*-values *X_v_* = Φ^−1^(1 − *p_v_*) are distributed as *N*(0,1) for genes satisfying the null hypothesis and *N*(*μ*, 1) for genes satisfying the alternative hypothesis [46]. Normal distributions also have been used to model transformed *p*-values from differential gene expression experiments [69, 61, 90].

More generally, the Normally Distributed Altered Subnetwork Problem is related to a larger class of *network anomaly* problems, which have been studied extensively in the machine learning and statistics literature [6, 4, 1, 3, 83, 82, 81, 80, 5]. To better understand the relationships between these problems and the algorithms developed to solve them, we will describe a generalization of the Altered Subnetwork Problem. We start by defining the following distribution, which generalizes the connected subnetworks in the Normally Distributed Altered Subnetwork Problem to any family of altered subsets.

#### Normally Distributed Altered Subset Distribution (ASD)

*Let n* > 0 *be a positive integer, let* 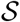 *be a family of subsets of* {1,…,*n*}, *and let* 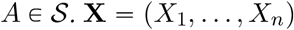 *is distributed according to the* Normally Distributed Altered Subset Distribution 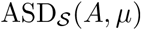 *provided*

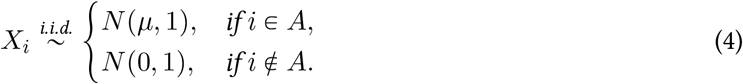

*Here, μ* > 0 *is the* mean *of the ASD and A is the* altered *subset of the ASD*.

More generally, the Altered Subset Distribution can be defined for any background distribution *D_B_* and altered distribution *D_A_*. We will restrict ourselves to normal distributions in accordance with the Normally Distributed Altered Subnetwork Problem, and we will subsequently assume normal distributions in both the Altered Subset Distribution and the Altered Subnetwork Problem.

The distribution in the Altered Subnetwork Problem is the 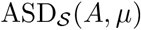, where the family 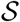 of subsets are connected subgraphs of the network *G*. In this terminology, the Altered Subnetwork Problem is the problem of estimating the parameters *A* and *μ* of the Altered Subset Distribution given data 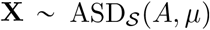 and knowledge of the parameter space 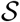 of altered subnetworks *A*. Thus, we generalize the Altered Subnetwork Problem to the ASD Estimation Problem, defined as follows.

#### ASD Estimation Problem

*Let* 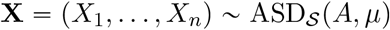. *Given* **X** *and* 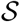, *find A and μ*.

The ASD Estimation Problem is a general problem of estimating the parameters of a *structured* alternative distribution. Different choices of 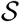 for the ASD Estimation Problem yield a number of interesting problems, some of which have been previously studied.

- 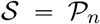, the power set of all subsets of {1,…,*n*}. We call the distribution 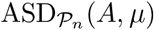 the *unstructured* ASD.
- 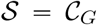, the set of all connected subgraphs of a graph *G* = (*V*, *E*). We call 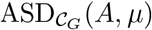 the *connected* ASD. The connected ASD Estimation Problem is equivalent to the Altered Subnetwork Problem described above.
- 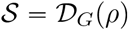, the set of all subgraphs of a graph *G* = (*V*, *E*) with edge density ⩾ *ρ*. [38, 88, 7] identify altered subnetworks with high edge density, and [2] identifies altered subnetworks with edge density *ρ* =1, i.e., cliques.
- 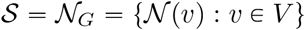, the set of all first-order network neighborhoods of a graph *G* = (*V*, *E*). [15, 44] use first-order network neighborhoods to prioritize cancer genes.
- 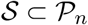, a family of subsets. Typically, 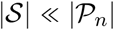 and 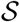 is not defined in terms of a graph. A classic example is gene set analysis; see [47] for a review.

### 2.2 Bias in Maximum Likelihood Estimation of the ASD

One reasonable approach for solving the ASD Estimation Problem is to compute a maximum likelihood estimator (MLE) for the parameters of the ASD. We derive the MLE below and show that it has undesirable statistical properties. All proofs are in the supplement.

#### Theorem 1.

*Let* 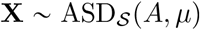. *The maximum likelihood estimators (MLEs) Â_ASD_ and* 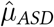 *of A and μ*, *respectively, are*

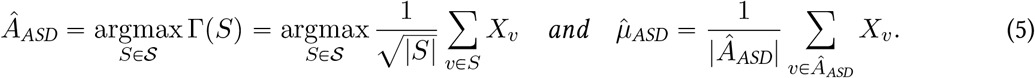

The maximization of Γ over 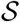 in (5) is a version of the *scan statistic*, a commonly used statistic to study point processes on lines and rectangles under various distributions [53, 34]. Comparing (5) and (2), we see that jActiveModules [48] computes the scan statistic over the family 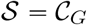 of connected subgraphs of the graph *G*. Thus, although jActiveModules [48] neither specifies the anomalous distribution *D_A_* nor provides a statistical justification for their subnetwork scoring function, Theorem 1 above shows that jActiveModules implicitly assumes that *D_A_* is a normal distribution, and that jActiveModules aims to solve the Altered Subnetwork Problem by finding the MLE *Â*_ASD_.

Despite this insight that jActiveModules computes the MLE, it has been observed that jActiveModules often identifies large subnetworks. [67] notes that the subnetworks identified by jActiveModules are large and “hard to interpret biologically”. They attribute the tendency of jActiveModules to identify large subnetworks to the fact that a graph typically has more large subnetworks than small ones. While this observation about the relative numbers of subnetworks of different sizes is correct, we argue that this tendency of jActiveModules to identify large subnetworks is due to a more fundamental reason: the MLE *Â*_ASD_ is a *biased* estimator of *A*.

First, we recall the definitions of bias and consistency for an estimator 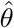 of a parameter *θ*.

#### Definition 1.

*Let* 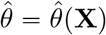 *be an estimator of a parameter θ given observed data* **X** = (*X*_1_,…,*X_n_*).(*a*) *The bias in the estimator* 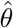 *of θ is* 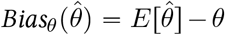. We say that 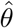 *is a biased estimator of θ if* 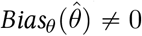, *and is an* unbiased *estimator of θ otherwise. (b) We say that* 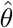 *is a* consistent *estimator of θ if* 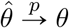, *where* 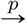 *denotes convergence in probability as n* → ∞, *and is an* inconsistent *estimator of θ otherwise*.

When it is clear from context, we omit the subscript *θ* and write 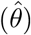 for the bias of estimator 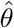.

Let 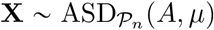 be distributed according to the unstructured ASD. We observe that the estimators |*Â*_ASD_|/*n* and 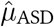 are both biased and inconsistent when both |*A*|/*n* and *μ* are moderately small (Figure 2). We summarize these observations in the following conjecture.

**Figure 2:**
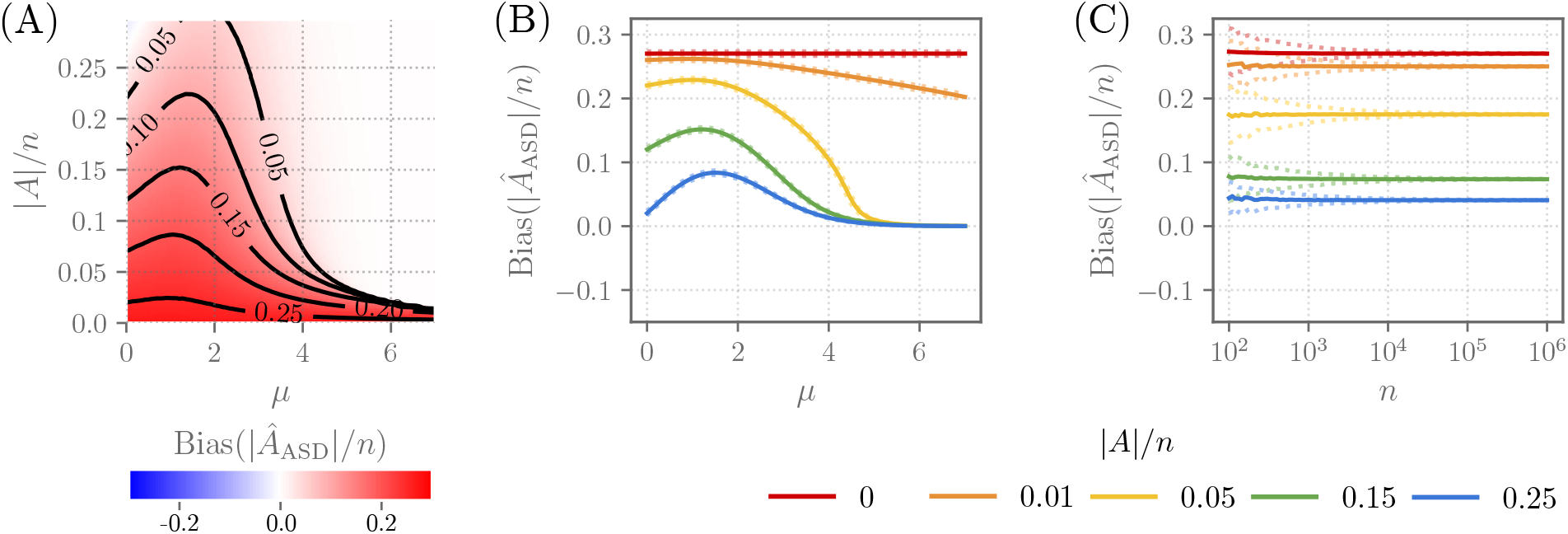
Scores 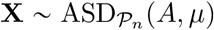 are distributed according to the unstructured ASD. (A) Bias(|*Â*_ASD_|/*n*) in the maximum likelihood estimate of |*A*|/*n* as a function of the mean *μ* and altered subset size |*A*|/*n* for *n* = 10^4^. (B) Bias(|*Â*_ASD_|/*n*) for *n* = 10^4^ and several values of |*A*|/*n*. Dotted lines indicate first and third quartiles in the estimate of the bias. (C) Bias (|*Â*_ASD_|/*n*) as a function of *n* for *μ* = 3 and for several values of |*A*|/*n*.

#### Conjecture.

*Let* 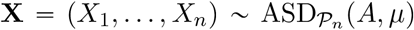. *Then there exist μ*_0_ > 0 *and β* > 0 *such that, if μ* < *μ*_0_ *and* |*A*|/*n* < *β*, *then* |*Â_ASD_*|/*n and* 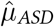 *are biased and inconsistent estimators of* |*A*|/*n and μ*, *respectively*.

Note that there are many examples in the literature of biased MLEs; e.g., the MLE for the variance of a (univariate) normal distribution or the MLE for the inverse of the mean of a Poisson distribution [30]. However, examples of inconsistent MLEs are somewhat rare [29].

Although we do not have a proof of the above conjecture, we prove the following results that partially explain the bias and inconsistency of the estimators |*A*_ASD_| and *μ*_ASD_. For the bias, we prove the following.

#### Theorem 2.

*Let* 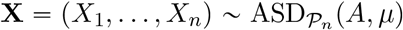 *with A* = ∅. *Then* |*Â_ASD_*| = *cn for sufficiently large n and with high probability, where* 0 < *c* < 0.35 *is independent of n*.

Empirically, we observe *c* ≈ 0.27, i.e., *Â_ASD_* contains more than a quarter of the scores (Figure 2). This closely aligns with the observation in [67] that jActiveModules reports subnetworks that contain approximately 29% of all nodes in the graph. Based on Theorem 2, one may suspect that |*Â*_ASD_| ≈ *cn* when *μ* or |*A*|/*n* is sufficiently small, providing some intuition for why |*Â*_ASD_|/*n* is biased. For inconsistency, we prove that the bias is independent of *n*.

#### Theorem 3.

*Let* 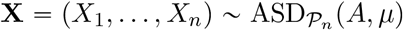, *where* |*A*| = *θ*(*n*). *For sufficiently large n*, *Bias*(|*Â_ASD_*|/*n*) *and* 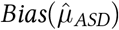 *are independent of n*.

## 3 The NetMix Algorithm

Following the observation that the maximum likelihood estimators of the distribution 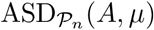 are biased, we aim to find a less biased estimator by explicitly modeling the distribution of the scores **X**. In this section, we derive a new algorithm, NetMix, that solves the Altered Subnetwork Problem by fitting a Gaussian mixture model (GMM) to **X**.

### 3.1 Gaussian Mixture Model

We start by recalling the definition of a GMM.

#### Gaussian Mixture Model

*Let μ* > 0 *and α* ∈ (0,1). *X is distributed according to the* Gaussian mixture model GMM(*α*, *μ*) *with parameters α and μ provided*

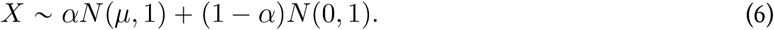

An alternate interpretation of the GMM is to draw a latent variable *Z* ~ Bernoulli(*α*) and select *X* ~ *N*(*μ*, 1) if *Z* = 1, and *X* ~ *N*(0,1) if *Z* = 0.

Given data **X** = (*X*_1_,…,*X_n_*), we define 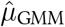 and 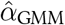 to be the MLEs for *μ* and *α*, respectively, obtained by fitting a GMM to **X**. In practice, 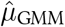 and 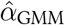 are obtained by the EM algorithm, which is known to converge to the MLEs as the number of samples goes to infinity [92, 23]. Furthermore, if 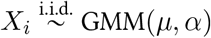 are distributed according to the GMM with *α* ≠ 0, then 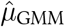 and 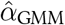 are consistent (and therefore asymptotically unbiased) estimators of *μ* and *α*, respectively [14].

Analogously, by fitting a GMM to data 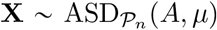 from the unstructured ASD, we observe that 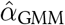 is a less biased estimator of |*A*|/*n* than |*Â*_ASD_|/*n* (Figure 3A,B). We also observe that 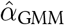 is a consistent estimator of |*A*|/*n* (Figure 3C). We summarize our findings in the following conjecture.

**Figure 3:**
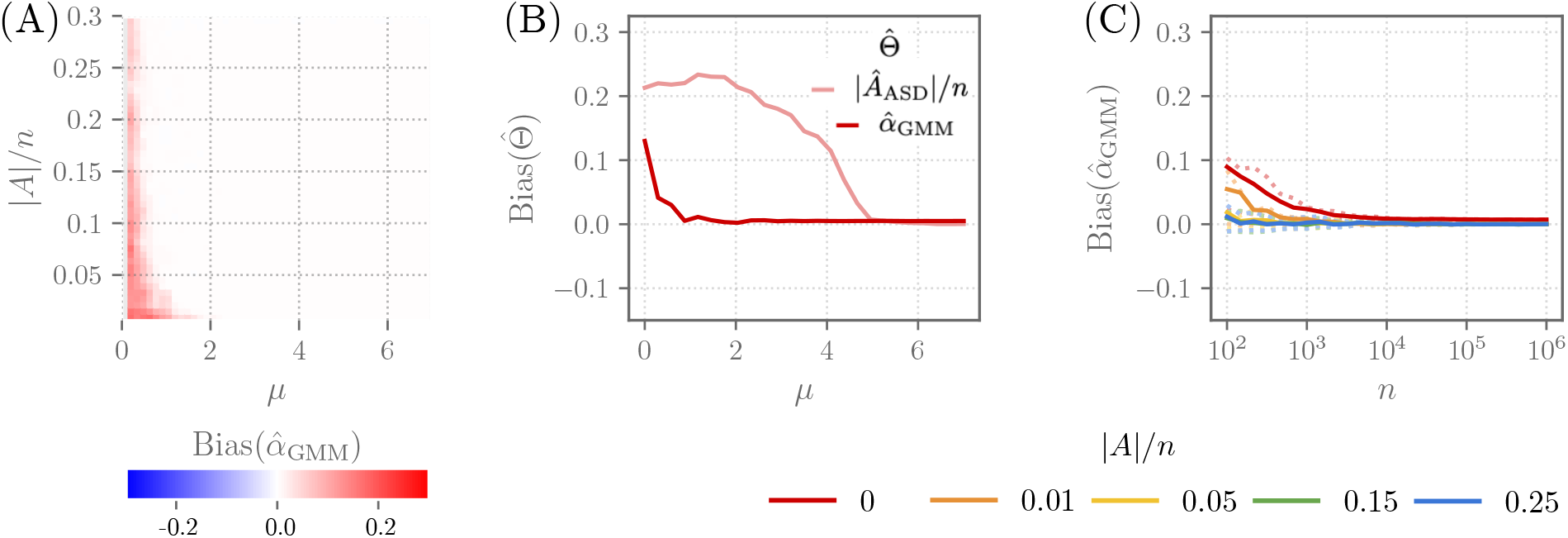
Scores 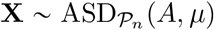 are distributed according to the unstructured ASD, and parameters 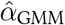 and 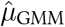 are obtained by the EM algorithm. (A) 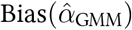 as a function of the mean *μ* and altered subnetwork size |*A*|/*n* for n = 10^4^. Compare with Figure 2A. (B) 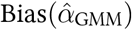 and Bias(|*Â*_ASD_|/*n*) as functions of the mean *μ* for |*A*|/*n* = 0.05 and *n* = 10^4^. (C) 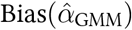 as a function of *n* for mean *μ* = 3 and several values of |*A*|/*n*. Compare with Figure 2C.

##### Conjecture.

*Let* 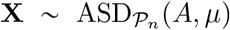 *with* |*A*| > 0, *and let Â*_ASD_ *be the MLE of A as defined in* (5). *Let* 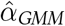 *and* 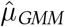 *be the MLEs of α and μ obtained by fitting a GMM to* **X**. *Then* 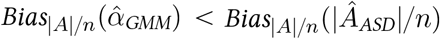. *Moreover*, 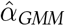 *and* 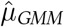 *are consistent estimators of* |*A*|/*n and μ, respectively*.

Although we do not have a proof of the above conjecture, a partial justification is the following relationship between the unstructured ASD and the GMM distribution. Let **X** = (*X*_1_,…,*X_n_*) be drawn from a mixture of unstructured ASDs over all possible anomalous sets *A* of size *k*, i.e., 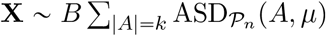, where 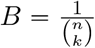 is a normalizing constant. Let 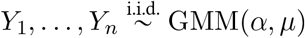 be independent samples from the GMM for *μ* > 0 and 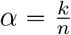 with corresponding latent variables *Z*_1_,…,*Z_n_*. Then, the joint distribution of the GMM samples **Y** = (*Y*_1_,…,*Y_n_*) conditioned on 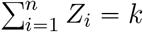 is equal to the distribution of **X**:

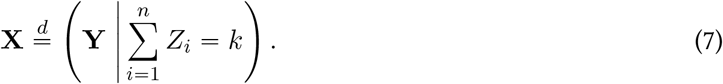

### 3.2 NetMix Algorithm

We derive an algorithm, NetMix, that uses the maximum likelihood estimators (MLEs) 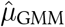 and 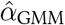 from the GMM to solve the Altered Subnetwork Problem. Note that the GMM is *not* identical to ASD, the distribution that generated the data. Despite this difference in distributions, the above conjecture provides justification that the GMM will yield less biased estimators of *A* and *μ* than the MLEs of the ASD distribution.

Given a graph *G* = (*V*, *E*) and scores **X** = (*X_v_*)_*v*∈*V*_, NetMix first computes the *responsibility r_v_* = Pr(*v* ∈ *A* | *X_v_*), or the probability that *v* ∈ *A*, for each vertex *v* ∈ *V*. The responsibilities *r_v_* are computed from the GMM MLEs 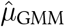 and 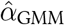 (which are estimated by the EM algorithm [24]) according to the formula

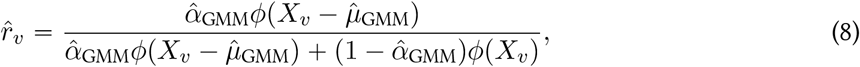

where *ϕ* is the PDF of the standard normal distribution.

Next, NetMix aims to find a connected subgraph *C* of size |*C*| ≈ *nα* that maximizes Σ_*v*∈*C*_ *r_v_*. In order to find such a subgraph, NetMix assigns a weight 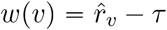 to each vertex *v*, where *τ* is chosen so that approximately 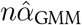 nodes have non-negative weights. NetMix then computes the maximum weight connected subgraph (MWCS) *Â*_NetMix_ in *G* by adapting the integer linear program in [27]. The use of *τ* is motivated by the observation that, if 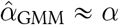, then we expect 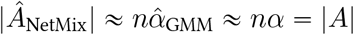.

We formally describe the NetMix algorithm for solving the Altered Subnetwork Problem below.

#### NetMix algorithm

Given a network *G* = (*V*, *E*) and vertex scores **X** = (*X_v_*)_*v*∈*V*_,

1. Compute 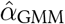 and 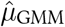, the MLEs of *α* and *μ*, by fitting a GMM to **X** using expectation maximization (EM).
2. Compute the estimated responsibilities 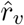 for each vertex *v* using (8).
3. Compute *τ* such that 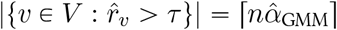, where ⌈·⌉ is the ceiling function.
4. Find the connected subgraph *Â*_NetMix_ defined by

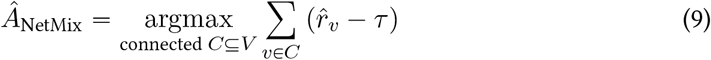

using integer linear programming.

NetMix bears some similarities to heinz [27], another algorithm to identify altered subnetworks. However, there are two important differences. First, heinz does not solve the Altered Subnetwork Problem defined in the previous section. Instead, heinz models the vertex scores (assumed to be *p*-values) with a beta-uniform mixture (BUM) distribution. The motivation for the BUM is based on an empirical goodness-of-fit in [72]; however, later work by the same author [71] observes that the BUM tends to underestimate the number of *p*-values drawn from the altered distribution. Second, heinz requires that the user specify a False Discovery Rate (FDR) and shifts the *p*-values according to this FDR. We show below that nearly all choices of the FDR lead to a biased estimate of |*A*|. Moreover, the manually selected FDR allows users to selectively tune the value of this parameter to influence which genes are in the inferred altered subnetwork, analogous to “*p*-hacking”[49, 68, 41]. Indeed, recently published analyses using heinz [17, 40, 52] use a wide range of FDR values. See the supplement for more details on the differences between heinz and NetMix. Despite these limitations, the ILP given in heinz to solve the MWCS problem is very useful for implementing NetMix and for computing the scan statistic (2) used in jActiveModules (see below).

## 4 Results

We compared NetMix to jActiveModules [48] and heinz [27] on simulated instances of the Altered Subnetwork Problem and on real datasets, including differential gene expression experiments from the Expression Atlas [70] and somatic mutations in cancer. jActiveModules is accessible only through Cytoscape [78, 19] and not a command-line interface, making it difficult to run on large number of a datasets. Thus, we implemented jActiveModules*, which computes the scan statistic (5) by adapting the integer linear program in heinz^2^. jActiveModules* output the global optimum of the scan statistic, while jActiveModules relies on heuristics (simulated annealing and greedy search) to find a local optimum.

### 4.1 Simulated Data

We compared NetMix, jActiveModules*, and heinz on simulated instances of the Altered Subnetwork Problem using the HINT+HI interaction network [57], a combination of binary and co-complex interactions in HINT [22] with high-throughput derived interactions from the HI network [76] as the graph *G*. For each instance, we randomly selected a connected subgraph *A* ⊆ *V* with size |*A*| = 0.05*n* using the random walk method of [59], and drew a sample 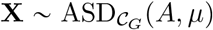. We ran each method on **X** and *G* to obtain an estimate *Â* of the altered subnetwork *A*. We ran heinz with three different choices of the FDR parameter (FDR = 0.001, FDR = 0.1, and FDR = 0.5) to reflect the variety of FDRs used in practice.

We found that NetMix output subnetworks whose size |*Â*_NetMix_| was very close to the true size across all values of *μ* in the simulations (Figure 4A). In contrast, jActiveModules* output subnetworks that were much larger than the implanted subnetwork for *μ* < 5. This behavior is consistent with our conjectures above about the large bias in the maximum likelihood estimator *Â*_ASD_ for the unstructured ASD. Note that *μ* > 5 corresponds to a large separation between the background and alternative distributions, and the network is not needed to separate these two distributions.

**Figure 4:**
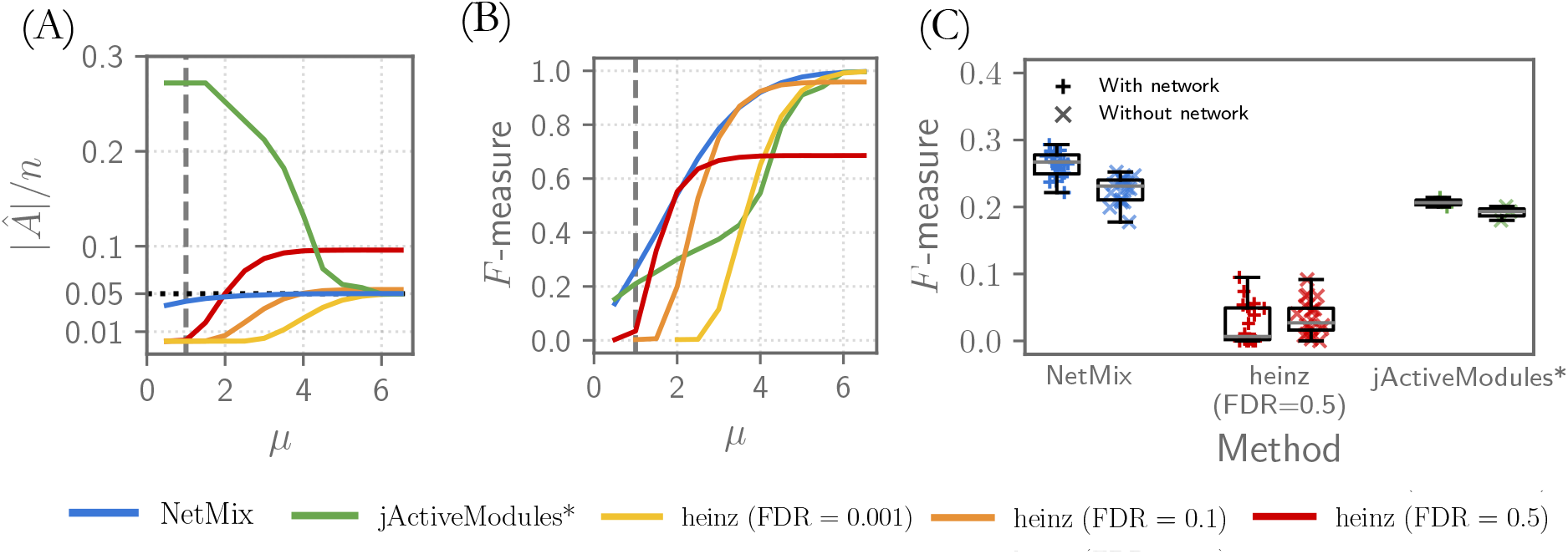
Comparison of altered subnetwork identification methods on simulated instances of the Altered Subnetwork Problem using the HINT+HI interaction network with *n* = 15074 nodes, and where the altered subnetwork *A* has size |*A*| = 0.05*n*. [Dashed vertical line (*μ* = 1) represents the smallest *μ* such that one can detect whether *G* contains an altered subnetwork^3^]. (A) Size |*Â*|/*n* of identified altered subnetwork *Â* as a function of mean *μ*. (B) *F*-measure for *Â* as a function of *μ*. (C) *F*-measure for *Â* at *μ* = 1, comparing performance with the network (left series for each method) and without the network (right series for each method).

We also quantified the overlap between the true altered subnetwork *A* and the subnetwork *Â* output by each method using the *F*-measure, finding that NetMix outperforms other methods across the full range of *μ* (Figure 4B). heinz requires the user to select an FDR value, and we find that the size of the output subnetwork and the *F*-measure varies considerably for different FDR (Figure 4A, 4B). When *μ* was small, a high FDR value (FDR = 0.5) yielded the best performance in terms of *F*-measure. However, when *μ* was large, a low FDR value (FDR = 0.001) gave better performance. While there are FDR values where the performance of heinz is similar to NetMix, the user *does not know what FDR value to select* for any given input, as the values of *μ* and the size |*A*| of the altered subnetwork are unknown.

The bias in |*Â*|/*n* observed using jActiveModules* with the interaction network (Figure 4A) was similar to the bias for the unstructured ASD (Figure 2A). Thus, we also evaluated how much benefit the network provided for each method. For small *μ*, we found that NetMix had a small but noticeable gain in performance when using the network; in contrast, other methods had nearly the same performance with or without the network (Figure 4C with further details in the supplement). These results emphasize the importance of evaluating network methods on simulated data *and* demonstrating that a network method outperforms a single-gene test; neither of these were done in the jActiveModules [48] and heinz [27] papers, nor are they common in many other papers on biological network analysis.

### 4.2 Differential Gene Expression Subnetworks

We compared NetMix, jActiveModules*, and heinz on gene expression data from the Expression Atlas [70]. We analyzed 945 differential expression experiments including 292 RNA-seq experiments and 653 microarray experiments. For 84% of these experiments, the GMM used by NetMix provided a better fit to the *p*-value distributions than the beta-uniform mixture (BUM) [72] used by heinz (see the supplement for more details). In addition, the GMM provided a better fit in 83/85 experiments where the null proportion (fraction of genes not differentially expressed) estimated by the GMM and BUM differed by ⩾ 0.25. In all 85 of these experiments, the BUM estimated a higher null proportion, consistent with the report in [71] that the BUM tends to overestimate the null proportion.

As many experiments had a small null proportion (i.e., most genes in the experiment were differentially expressed), we restricted our analysis to the 157 experiments from the Expression Atlas with a null proportion ⩾ 0.8 as estimated by the GMM. We ran NetMix, jActiveModules*, and heinz on these 157 experiments with the HINT+HI network. For heinz, we used three FDR values: FDR = 0.1, FDR = 0.001, and the FDR value such that |*Â*_NetMix_| genes have a positive weight in the heinz scoring. These choices demonstrate how users might “*p*-hack” the FDR value to achieve desirable results. We also compared to a method that ignores network topology, selecting the |*Â*_NetMix_| genes with the lowest *p*-values; we call this method “top *p*-values”. See the supplement for specific details on these methods.

Both NetMix and heinz identified subnetworks that were significantly smaller than jActiveModules* (Figure 5A), which is consistent with previous observations [67] that jActiveModules estimates overly large subnetworks. At the same time, NetMix identified subnetworks with significant overlap (FDR ⩽ 0.01) with more biological process GO terms than heinz (*p* = 3.3 · 10^−12^, *t*-test; Figure 5B) or top *p*-values (*p* < 2.2 · 10^−16^, *t*-test; Figure 5B). We also found that subnetworks identified by NetMix had greater overlap (as quantified by *F*-measure) with genes in the top *k* GO terms (Figure 5C). These results suggest that NetMix identifies subnetworks that are more relevant to differential expression experiments than other methods.

**Figure 5:**
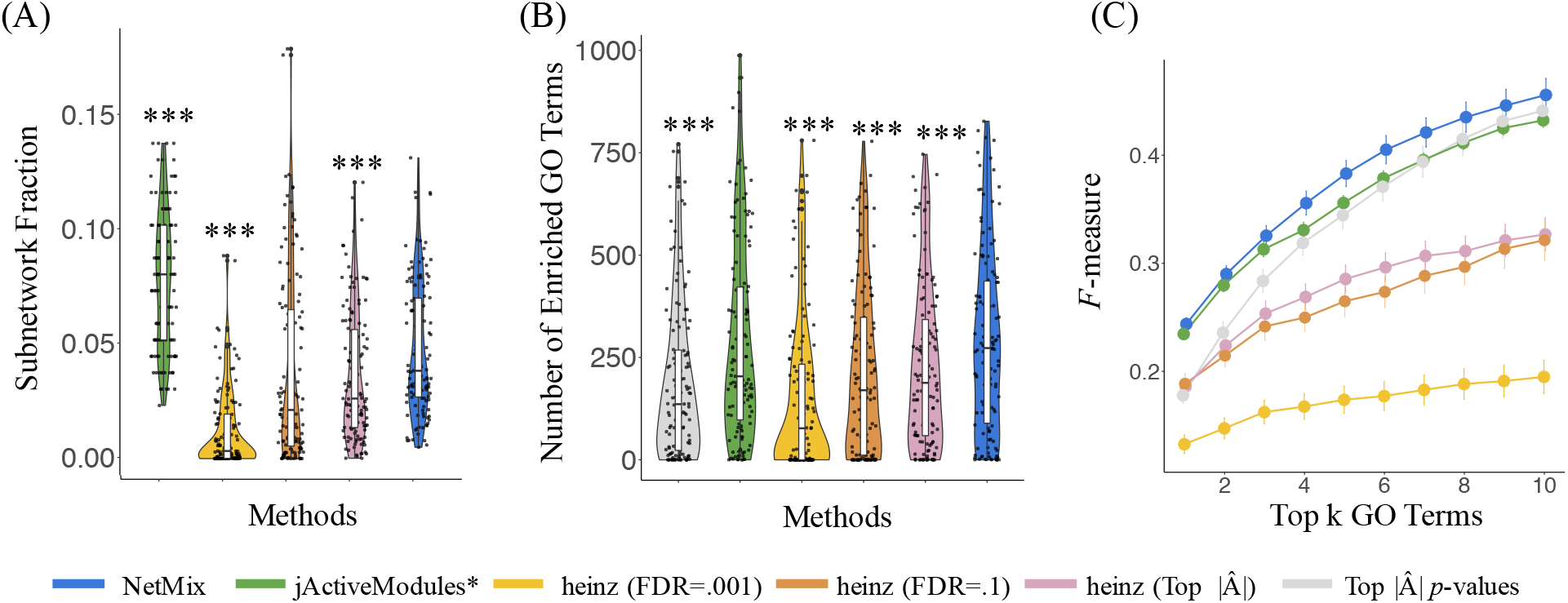
(A) Fraction of genes in the HINT+HI interaction network that are in the subnetwork identified by each method. *: *p* ⩽ 0.01, ≈: *p* ⩽ 0.001, * * *: *p* ⩽ 10^−4^ indicate significant *p*-values in paired *t*-test between NetMix and other methods. (B) Number of enriched GO biological process terms for altered subsets identified by each method. (C) *F*-measure of the *k* most enriched GO terms.

We examined the experiment E-GEOD-11199 in more detail. This experiment compared *Mtb*-stimulated and unstimulated macrophages [86]. NetMix identified a subnetwork containing 706 genes, half the size of the jActiveModules* subnetwork containing 1450 genes. Both of these subnetworks contained 37 of the 42 genes whose differential expression was experimentally validated by RT-PCR [86]. Although the NetMix subnetwork was less than half the size of the jActiveModules* subnetwork, the NetMix subnetwork overlapped more GO terms (445 vs. 179). In contrast, heinz (using FDR = 0.27) identified a subnetwork of 382 genes containing only 25 RT-PCR validated genes. Finally, the 692 genes with the smallest *p*-values include only 7 validated genes. These results show that the NetMix subnetwork contains many biologically relevant genes, including most of the RT-PCR validated genes, without being overly large.

### 4.3 Somatic Mutations In Cancer

We compared the performance of NetMix, jActiveModules* [4, 1], jActiveModules [48], heinz [27] and Hierarchical HotNet [75] in identifying cancer driver genes, using the MutSig2CV driver *p*-values [54] from the TCGA PanCanAtlas project [8]. We ran all methods on the HINT+HI interaction network described above, as well as the iRefIndex 15.0 [74] and ReactomeFI 2016 [21, 28] interaction networks. See the supplement for more details on the datasets.

We evaluated the quality of the subnetwork *Â* reported by each method by computing the overlap with the list of cancer genes in the COSMIC Cancer Gene Census (CGC) [33, 31]. We found that NetMix outperforms all other methods in *F*-measure across all interaction networks. For example, using the HINT+HI network, NetMix achieved an *F*-measure of 0.277, compared to *F*-measures of 0.191 for jActiveModules*, 0.216 for heinz (FDR = 0.001), 0.264 for heinz (FDR = 0.1), and 0.214 for Hierarchical HotNet^4^. Both the NetMix and Hierarchical HotNet results were statistically significant (*p* < 0.01) on all 3 interaction networks according to permutation tests from [75]. The modest *F*-measures for all methods are not surprising; the genes in CGC have diverse alterations across cancer types and thus high recall is not expected by this restricted analysis of single-nucleotide mutations in a subset of cancer types. Nevertheless, the higher performance of NetMix on this task across all networks is encouraging. Further details of these comparisons are in the supplement.

## 5 Discussion

In this paper, we revisit the classic problem of identifying altered subnetworks in high-throughput biological data. We formalize the Altered Subnetwork Problem as the estimation of the parameters of the Altered Subset Distribution (ASD). We show that the seminal algorithm for this problem, jActiveModules [48], is equivalent to a maximum likelihood estimator (MLE) of the ASD. At the same time, we show that the MLE is a biased estimator of the altered subnetwork, with especially large positive bias for small altered subnetworks. This bias explains previous reports that jActiveModules tends to output large subnetworks [67].

We leverage these observations to design NetMix, a new algorithm for the Altered Subnetwork Problem. We show that NetMix outperforms existing methods on simulated and real data. NetMix fits a Gaussian mixture model (GMM) to observed node scores and then finds a maximum weighted connected subgraph using node weights derived from the GMM. heinz [27], another widely used method for altered subnetwork identification, also derives node weights from a mixture model (a beta-uniform mixture of *p*-values) and finds a maximum weighted connected subgraph. However, heinz does not solve the Altered Subnetwork Problem in a strict sense; rather, heinz requires users to choose a parameter (an FDR estimate for the mixture fit) that implicitly constrains the size of the identified subnetwork. This user-defined parameter encourages *p*-hacking [49, 68, 41], and we find that nearly all values of this parameter lead to biased estimates of the size of the altered subnetwork.

We note a number of directions for future work. The first is to generalize our theoretical contributions to the identification of *multiple* altered subnetworks, a situation which is common in biological applications where multiple biological processes may be perturbed [62]. While it is straightforward to run NetMix iteratively to identify multiple subnetworks – as jActiveModules does – a rigorous assessment of the identification of multiple altered subnetworks would be of interest. Second, our results on simulated data (Section 4.1) show that altered subnetwork methods have only marginal gains over simpler methods that rank vertices without information from network interactions. We hypothesize that this is because connectivity is not a strong constraint for biological networks; indeed the biological interaction networks that we use have both small diameter and small average shortest path between nodes (see the supplement for specific statistics). Specifically, we suspect that most subsets of nodes are “close” to a connected subnetwork in such biological networks, and thus the MLE of connected altered subnetworks has similar bias as the MLE of the unstructured altered subset distribution. In contrast, for other network topologies like the line graph, connectivity is a much stronger topological constraint (see the supplement for a brief comparison of different topologies). It would be useful to investigate this hypothesis and characterize the conditions when networks provide benefit for finding altered subnetworks. In particular, other topological constraints such as dense subgraphs [38, 7], cliques [2], and subgraphs resulting from heat diffusion and network propagation processes [87, 88, 57, 20] have been used used to model altered subnetworks in biological data. Generalizing the theoretical results in this paper to these other topological constraints may be helpful for understanding the parameter regimes where these topological constraints provide signal for identification of altered subnetorks. Finally, we note that biological networks often have substantial ascertainment bias, with more interactions annotated for well-studied genes [44, 76], and these well-studied genes in turn may also be more likely to have outlier measurements/scores. Thus, any network method should carefully quantify the regime where it outperforms straightforward approaches – e.g., methods based on ranking nodes by gene scores or node degree – both on well-calibrated simulations and on real data.

## Acknowledgments

We thank Mohammed El-Kebir for assistance with implementing jActiveModules* by modifying the ILP in heinz. We thank David Tse for directing us to the network anomaly literature. M.A.R. was supported in part by the National Cancer Institute of the NIH (Cancer Target Discovery and Development Network grant U01CA217875). B.J.R. was supported by US National Institutes of Health (NIH) grants R01HG007069 and U24CA211000.

## Footnotes

1 jActiveModules actually maximizes Γ_norm_(*S*) = (Γ(*S*) − *μ*_|*S*|_)/*σ*_|*S*|_, a *z*-score normalized version of Γ(*S*), where *μ*_|*S*|_ and *σ*_|*S*|_ are the mean and standard deviation, respectively, of Γ(*T*) over all subsets *T* ⊆ *V* of size |*S*|. We show in the supplement that maximizing Γ_norm_(*S*) is equivalent to maximizing the unnormalized Γ(*S*) when the data is generated from normal distributions.

2 The scan statistic (2) is the maximization of a non-linear objective function, but for fixed subnetwork size |*S*| the objective function is linear. We computed the scan statistic by modifying the ILP in heinz [27] and running this ILP over all possible subnetwork sizes.

3 Formally, *μ* is the smallest mean such that the hypotheses 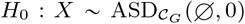 and 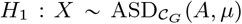 are asymptotically distinguishable. See [83] for details.

4 The jActiveModules greedy search algorithm failed to complete within 100 hours, while the jActiveModules simulated annealing algorithm yielded an *F*-measure of 0:086

