## Supplementary Material for "NetMix: A network-structured mixture model for reduced-bias estimation of altered subnetworks"

The following sections elaborate on the methods, data, and results from the main manuscript.

#### 1 Altered Subnetworks, Altered Subsets, and Maximum Likelihood Estimation

##### 1.1 $z$-score normalized scan statistic

Let $\mathbf{X} = (X_1, \dots, X_n) \sim \text{ASD}_{\mathcal{S}}(A, \mu)$. Define $\mu_k$ and $\sigma_k$ to be the mean and standard deviation, respectively, of $\Gamma(T)$ over all subsets $T \in \mathcal{S}$ with $|T| = k$. Define $\Gamma_{\text{norm}}(S) = \frac{\Gamma(S) - \mu_{|S|}}{\sigma_{|S|}}$, i.e. $\Gamma_{\text{norm}}(S)$ is the $z$-score normalized scan statistic that is implemented in the jActiveModules algorithm [12].

We show below that the $z$-score normalization of the scan statistic does not affect the maximum likelihood estimates of $A$ and $\mu$ when $\mathbf{X} \sim \text{ASD}_{\mathcal{P}_n}(A, \mu)$ is distributed according to the unstructured ASD.

**Lemma 1.** *Let $\mathbf{X} \sim \text{ASD}_{\mathcal{P}_n}(A, \mu)$ be distributed according to the unstructured ASD. Then*

$$\hat{A}_{ASD} = \underset{S \in \mathcal{P}_n}{\text{argmax}}\, \Gamma_{norm}(S). \tag{1}$$

*Proof.* For any $k > 0$, we have

$$\begin{aligned} \mu_k &= E\left[\Gamma(T) \mid |T| = k\right] \\ &= E\left[\frac{1}{|T|}\sum_{v \in T} X_v \,\middle|\, |T| = k\right] \\ &= \frac{1}{k}\left(\frac{|A|}{n} \cdot k \cdot \mu + \left(1 - \frac{|A|}{n}\right) k \cdot 0\right) \\ &= \frac{|A|}{n}\mu, \end{aligned} \tag{2}$$

where we use the fact that, for a random subset $T \in \mathcal{P}_n$ of size $|T| = k$, then $\frac{|A|}{n}$ fraction of elements in $T$ are distributed as $N(\mu, 1)$ and $1 - \frac{|A|}{n}$ fraction of elements in $T$ are distributed as $N(0, 1)$. This follows from the formula for the mean of a hypergeometric distribution.

* These authors contributed equally.
**Contact:**

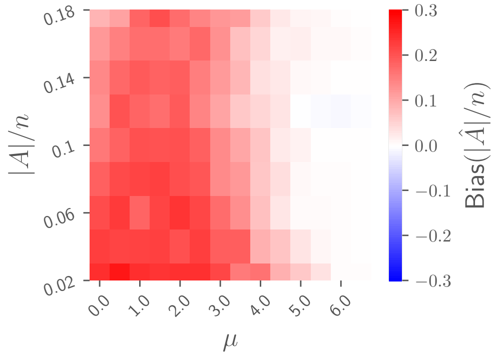


Figure S1: $\text{Bias}(\hat{A}_{\text{norm}}/n)$ for $n = 10^2$ as a function of the mean $\mu$ and the altered subset size $|A|/n$. Note the similarities to Figure 2a in the main text.

We also have that $\sigma_k = 1$ for all $k$, since the standard deviation of each $X_v$ is 1. Thus,

$$\Gamma_{\text{norm}}(S) = \frac{\Gamma(S) - \mu_{|S|}}{\sigma_{|S|}} = \Gamma(S) - \frac{|A|}{n}\mu. \tag{3}$$

Since $\mu \cdot (|A|/n)$ is independent of $S$, it follows that $\text{argmax}_{S \in \mathcal{S}} \Gamma_{\text{norm}}(S) = \text{argmax}_{S \in \mathcal{S}} \Gamma(S)$. □

We demonstrate Lemma 1 empirically in Figure S1.

#### 1.2 Proof of Theorem 1

**Theorem 1.** *Let* $\mathbf{X} \sim \text{ASD}_{\mathcal{S}}(A, \mu)$. *The maximum likelihood estimators (MLEs)* $\hat{A}_{ASD}$ *and* $\hat{\mu}_{ASD}$ *of* $A$ *and* $\mu$, *respectively, are*

$$\hat{A}_{ASD} = \underset{S \in \mathcal{S}}{\text{argmax}}\, \Gamma(S) = \underset{S \in \mathcal{S}}{\text{argmax}} \frac{1}{\sqrt{|S|}} \sum_{v \in S} X_v \quad \textit{and} \quad \hat{\mu}_{ASD} = \frac{1}{|\hat{A}_{ASD}|} \sum_{v \in \hat{A}_{ASD}} X_v. \tag{4}$$

*Proof.* The maximum likelihood estimates $\hat{A}_{\text{ASD}}$ and $\hat{\mu}_{\text{ASD}}$ are

$$\hat{A}_{\text{ASD}}, \hat{\mu}_{\text{ASD}} = \max_{S \in \mathcal{S}, \nu > 0} \text{L}(S, \nu \mid \mathbf{X})$$

where the likelihood function $\text{L}(S, \nu \mid \mathbf{X})$ is

$$\text{L}(S, \nu \mid \mathbf{X}) = \left( \prod_{v \in S} \frac{1}{\sqrt{2\pi}} \exp\left( -\frac{(X_v - \nu)^2}{2} \right) \right) \left( \prod_{v \notin S} \frac{1}{\sqrt{2\pi}} \exp\left( -\frac{X_v^2}{2} \right) \right). \tag{5}$$

Simplifying (5) and using the monotonicity of the logarithm:

$$\hat{A}_{\text{ASD}}, \hat{\mu}_{\text{ASD}} = \underset{A \in \mathcal{S},\, \mu > 0}{\text{argmin}} \left( \sum_{v \in A} (X_v - \mu)^2 + \sum_{v \notin A} X_v^2 \right). \tag{6}$$

Define $\bar{X}_A = \frac{1}{|A|}\sum_{v\in A} X_v$. Then, the value of $\mu$ that minimizes the right hand of (6) is $\mu = \bar{X}_A$. Plugging $\mu = \bar{X}_A$ into the right side of (6) and simplifying gives us:

$$\begin{aligned}
\sum_{v\in A}(X_v-\mu)^2 + \sum_{v\notin A} X_v^2 &= \sum_{v\in A}(X_v-\bar{X}_A)^2 + \sum_{v\notin A} X_v^2 \\
&= \left(\sum_{v\in A} X_v^2 - 2\bar{X}_A\left(\sum_{v\in A} X_v\right) + |A|\cdot\bar{X}_A^2\right) + \sum_{v\notin A} X_v^2 \\
&= \left(\sum_{v\in V} X_v^2\right) - 2|A|\cdot\bar{X}_A^2 + |A|\cdot\bar{X}_A^2 \qquad (7) \\
&= \left(\sum_{v\in V} X_v^2\right) - |A|\cdot\bar{X}_A^2 \\
&= \left(\sum_{v\in V} X_v^2\right) - \frac{1}{|A|}\left(\sum_{v\in A} X_v\right)^2.
\end{aligned}$$

Thus, we have

$$\hat{A}_{\text{ASD}} = \underset{A\in\mathcal{S}}{\operatorname{argmin}}\left(\left(\sum_{v\in V} X_v^2\right) - \frac{1}{|A|}\left(\sum_{v\in A} X_v\right)^2\right) \tag{8}$$

Since the first term is independent of $A$, we can simplify the above expression as

$$\hat{A}_{\text{ASD}} = \underset{A\in\mathcal{S}}{\operatorname{argmax}}\left(\frac{1}{|A|}\left(\sum_{v\in A} X_v\right)^2\right) = \underset{A\in\mathcal{S}}{\operatorname{argmax}}\left(\frac{1}{\sqrt{|A|}}\left(\sum_{v\in A} X_v\right)\right). \tag{9}$$

Finally, because the optimal $\mu$ is $\mu = \bar{X}_A$, we have

$$\hat{\mu}_{\text{ASD}} = \frac{1}{|\hat{A}_{\text{ASD}}|}\sum_{v\in\hat{A}_{\text{ASD}}} X_v. \qquad \square$$

##### 1.3 Proof of Theorem 2

Let $X_{[1]} \geqslant X_{[2]} \geqslant \cdots \geqslant X_{[n]}$ be the order statistics of $(X_1,\ldots,X_n)$. Note that we can rewrite $\hat{A}_{\text{ASD}}$ as

$$\hat{A}_{\text{ASD}} = \underset{A\in\mathcal{P}_n}{\operatorname{argmax}}\frac{1}{\sqrt{|A|}}\sum_{v\in\{1,\ldots,n\}} = \underset{k>0}{\operatorname{argmax}}\frac{1}{\sqrt{k}}\sum_{i=1}^{k} X_{[i]}, \tag{10}$$

To prove Theorem 2, we require the following high-probability bound on $\frac{1}{\sqrt{k}}\sum_{i=1}^{k} X_{[i]}$.

**Lemma 2.** *Let $X_i \overset{i.i.d.}{\sim} N(0,1)$, and let $Y_k = \frac{1}{\sqrt{k}}\sum_{i=1}^{k} X_{[i]}$. Then*

$$Y_k \leqslant \sqrt{2\log\binom{n}{k}} \tag{11}$$

*with high probability.*

*Proof.* Let $t = \sqrt{2\log\binom{n}{k}}$, and define $\mathcal{N}_k = \{B \in [n] : |B| = k\}$. We have

$$\begin{aligned} P(Y_k > t) &= P\left(\max_{B\in\mathcal{N}_k} \sum_{v\in B} X_v > t\sqrt{k}\right) \\ &\leqslant \sum_{B\in\mathcal{N}_k} P\left(\sum_{v\in B} X_v > t\sqrt{k}\right) \\ &= |\mathcal{N}_k| \cdot (1-\Phi(t)) \\ &= \binom{n}{k} \cdot (1-\Phi(t)), \end{aligned} \tag{12}$$

where the first inequality uses a union bound and the second equality uses that $\sum_{v\in B} X_v \sim N(0,k)$. Plugging in the well-known bound $1-\Phi(x) \leqslant \frac{1}{\sqrt{2\pi}}\frac{1}{x}e^{-x^2/2}$:

$$\begin{aligned} P(Y_k > t) &\leqslant \binom{n}{k} \cdot (1-\Phi(t)) \\ &\leqslant \binom{n}{k} \cdot \frac{1}{\sqrt{2\pi}}\frac{1}{t}e^{-\frac{t^2}{2}} \\ &= \binom{n}{k} \cdot \frac{1}{\sqrt{2\pi}}\frac{1}{\sqrt{2\log\binom{n}{k}}}e^{-\log\binom{n}{k}} \\ &= \frac{1}{\sqrt{2\pi}}\frac{1}{\sqrt{2\log\binom{n}{k}}} \end{aligned} \tag{13}$$

Because $\frac{1}{\sqrt{2\pi}}\frac{1}{\sqrt{2\log\binom{n}{k}}} \to 0$ as $n \to \infty$, it follows that $Y_k \leqslant t = \sqrt{2\log\binom{n}{k}}$ with high probability. □

We now prove Theorem 2.

**Theorem 2.** *Let* $\mathbf{X} = (X_1, \ldots, X_n) \sim \mathrm{ASD}_{\mathcal{P}_n}(A, \mu)$ *with* $A = \varnothing$. *Then* $|\hat{A}_{ASD}| = cn$ *for sufficiently large* $n$ *and with high probability, where* $0 < c < 0.35$ *is independent of* $n$.

*Proof.* We first prove that $|\hat{A}_{\mathrm{ASD}}| = \theta(n)$ with high probability.

Let $X_{[1]} \geqslant X_{[2]} \geqslant \cdots \geqslant X_{[n]}$ be the order statistics of $(X_1, \ldots, X_n)$. Let $Y_k = \frac{1}{\sqrt{k}}\sum_{i=1}^{k} X_{[i]}$. By (10), it suffices to show that $\mathrm{argmax}_k\, Y_k = \theta(n)$ with high probability.

By Lemma 2, we have $Y_k < \sqrt{2\log\binom{n}{k}}$ for all $k$ with high probability, and so by [9], it follows that $Y_k = o(\sqrt{n})$ for all $k = o(n)$ with high probability.

On the other hand, we have $E[X_{[n/4]}] = c$ for a fixed constant $c > 1$ and $\mathrm{Var}\left(X_{[n/4]}\right) = O\left(\frac{1}{k}\right)$ by [3]. So by Chebyshev's inequality, $X_{[n/4]} > 1$ with high probability, and it follows that

$$\begin{aligned} Y_{n/4} &= \frac{1}{\sqrt{\frac{n}{4}}}\sum_{i=1}^{\frac{n}{4}} X_{[k]} \\ &\geqslant \frac{1}{\sqrt{\frac{n}{4}}}\sum_{i=1}^{\frac{n}{4}} 1 \\ &\geqslant \sqrt{\frac{n}{4}} = \frac{\sqrt{n}}{2}. \end{aligned} \tag{14}$$

Thus,

$$Y_k = o(\sqrt{n}) < \frac{\sqrt{n}}{2} = Y_{n/4} \tag{15}$$

for all $k = o(n)$, and it follows that $\operatorname{argmax}_k Y_k = \theta(n)$.

Next, we show that $\operatorname{argmax}_k Y_k = cn$ for $0 < c < 0.35$. It is sufficient to show that $Y_k$ is decreasing for $k > 0.35n$ with high probability. Let $S_k = \sum_{i=1}^{k} X_{[i]}$. Note that $Y_k$ is decreasing iff

$$\begin{aligned} Y_k > Y_{k+1} &\Leftrightarrow \frac{1}{\sqrt{k}} \sum_{i=1}^{k} X_{[i]} > \frac{1}{\sqrt{k+1}} \sum_{i=1}^{k+1} X_{[i]} \\ &\Leftrightarrow X_{[k+1]} < S_k \left( \sqrt{1 + \frac{1}{k}} - 1 \right) \end{aligned} \tag{16}$$

For sufficiently large $k$, we have $\sqrt{1 + \frac{1}{k}} - 1 > \frac{1}{2.001k}$ by a Taylor series expansion, and $S_k \geqslant k\sqrt{\frac{2}{\pi}}$ by [9]. Moreover, for sufficiently large $n$, $X_{[0.35n]} < E[X_{[0.35n]}] + 0.0001 = 0.3981$ with high probability by Chebyshev's inequality. So for all $k \geqslant 0.35n$, we have

$$\begin{aligned} X_{[k+1]} &< X_{[0.35n]} \\ &< 0.3981 \\ &< \frac{1}{2.001}\sqrt{\frac{2}{\pi}} \\ &< \left( \sqrt{1 + \frac{1}{k}} - 1 \right) S_k. \end{aligned} \tag{17}$$

with high probability. ☐

#### 1.4 Proof of Theorem 3

**Theorem 3.** *Let* $\mathbf{X} = (X_1, \ldots, X_n) \sim \mathrm{ASD}_{\mathcal{P}_n}(A, \mu)$, *where* $|A| = \theta(n)$. *For large* $n$, *Bias*$(|\hat{A}_{ASD}|/n)$ *and Bias*$(\hat{\mu}_{ASD})$ *are independent of* $n$.

*Proof.* From Theorem 1, we have

$$\hat{A}_{\text{ASD}} = \max_{S \subset \{1,\ldots,n\}} \frac{1}{\sqrt{|S|}} \sum_{v \in S} X_v. \tag{18}$$

Because the maximum is taken over all subsets of $\{1, \ldots, n\}$, an equivalent formulation of the above is $\hat{A}_{\text{ASD}} = \{v : X_v > \hat{T}_{\text{ASD}}\}$, where

$$\hat{T}_{\text{ASD}} = \max_{T \in \mathbb{R}} \left( \frac{1}{\sqrt{\#\{v : X_v > T\}}} \sum_{v : X_v > T} X_v \right). \tag{19}$$

Now for any $T > 0$, we have that

$$\lim_{n \to \infty} \frac{\#\{v : X_v > T\}}{(1-\alpha)n \cdot (1 - \Phi(T)) + \alpha n \cdot (1 - \Phi(T - \mu))} = 1, \tag{20}$$

where $\Phi$ is the CDF of a standard normal. This is because, asymptotically, for the $(1-\alpha)n$ of the samples that are distributed as $N(0, 1)$, there is a $1 - \Phi(T)$ probability that an individual sample is greater than $T$. Similarly, asymptotically, for the $\alpha n$ samples that are distributed as $N(\mu, 1)$, there is a $1-\Phi(T-\mu)$ probability that an individual sample is greater than $T$.

Next, we compute the limit of $\sum_{v:X_v>T} X_v$ by dividing this sum into two parts:

$$\sum_{X_v>T} X_v = \sum_{X_v>T, v\notin A} X_v + \sum_{X_v>T, v\in A} X_v. \tag{21}$$

For the first sum, we note the following two equations:

$$\lim_{n\to\infty} \frac{|\{v : X_v > T, v \notin A\}|}{(1-\alpha)n \cdot (1-\Phi(T))} = 1 \tag{22}$$

$$\lim_{n\to\infty} \frac{\frac{1}{|\{v:X_v>T, v\notin A\}|} \sum_{X_v>T, v\notin A} X_v}{\eta(0,T)} = 1 \tag{23}$$

where $\eta(\nu, T)$ is the mean of a $N(\nu, 1)$ distributed truncated to lie in $(T, \infty)$. Thus, we have

$$\frac{\sum_{X_v>T, v\in A} X_v}{(1-\alpha)n \cdot (1-\Phi(T)) \cdot \eta(0,T)} = 1. \tag{24}$$

By similar logic for $\sum_{X_v>T, v\notin A} X_v$, we have

$$\lim_{n\to\infty} \frac{\sum_{X_v>T, v\in A} X_v}{\alpha n \cdot (1-\Phi(T-\mu)) \cdot \eta(\mu, T)} = 1. \tag{25}$$

Combining (24) and (25) yields

$$\lim_{n\to\infty} \frac{\sum_{X_v>T} X_v}{(1-\alpha)n(1-\Phi(T)) \cdot \eta(0,T) + \alpha n(1-\Phi(T-\mu)) \cdot \eta(\mu,T)} = 1, \tag{26}$$

and thus combining (20) with (26) gives us

$$\begin{aligned}
\lim_{n\to\infty} \hat{T}_{\text{ASD}} &= \lim_{n\to\infty} \left( \operatorname*{argmax}_{T\in\mathbb{R}} \frac{(1-\alpha)n(1-\Phi(T)) \cdot \eta(0,T) + \alpha n(1-\Phi(T-\mu)) \cdot \eta(\mu,T)}{\sqrt{(1-\alpha)n \cdot (1-\Phi(T)) + \alpha n \cdot (1-\Phi(T-\mu))}} \right) \\
&= \lim_{n\to\infty} \left( \operatorname*{argmax}_{T\in\mathbb{R}} \frac{(1-\alpha)(1-\Phi(T)) \cdot \eta(0,T) + \alpha(1-\Phi(T-\mu)) \cdot \eta(\mu,T)}{\sqrt{(1-\alpha) \cdot (1-\Phi(T)) + \alpha \cdot (1-\Phi(T-\mu))}} \cdot \sqrt{n} \right) \\
&= \operatorname*{argmax}_{T\in\mathbb{R}} \frac{(1-\alpha)(1-\Phi(T)) \cdot \eta(0,T) + \alpha(1-\Phi(T-\mu)) \cdot \eta(\mu,T)}{\sqrt{(1-\alpha) \cdot (1-\Phi(T)) + \alpha \cdot (1-\Phi(T-\mu))}}.
\end{aligned} \tag{27}$$

It follows that for sufficiently large $n$, $\hat{T}_{\text{ASD}}$ is independent of $n$.

Moreover, we have that $|\hat{A}_{\text{ASD}}| = \#\{v : X_v > \hat{T}_{\text{ASD}}\}$, so

$$E[|\hat{A}_{\text{ASD}}|/n] = (1-\alpha) \cdot (1-\Phi(\hat{T}_{\text{ASD}})) + \alpha \cdot (1-\Phi(\hat{T}_{\text{ASD}} - \mu)). \tag{28}$$

Since $\hat{T}_{\text{ASD}}$ is independent of $n$, it follows that $E[|\hat{A}_{\text{ASD}}|/n]$ is also independent of $n$. Thus, $\text{Bias}(|\hat{A}_{\text{ASD}}|/n) = E[|\hat{A}_{\text{ASD}}|/n] - |A|/n$ is independent of $n$. A similar calculation shows that $\text{Bias}(\hat{\mu}_{\text{ASD}})$ is also independent of $n$. □

### 2 The NetMix Algorithm

#### 2.1 Choice of $\tau$ in NetMix

In the NetMix algorithm, we choose $\tau = \tau_{\text{NetMix}}$ such that $|\{v \in V : \hat{r}_v - \tau > 0\}| = \lceil \hat{\alpha}_{\text{GMM}} n \rceil$. Another reasonable choice is $\tau = 0.5$, which corresponds to assigning non-negative weight $w(v) > 0$ to vertices with responsibility $\hat{r}_v \approx P(v \in A \mid X_v)$ greater than 0.5.

When using NetMix to solve the unstructured ASD estimation problem (i.e. the argmax in Step 4 of the NetMix algorithm is taken over all subsets $C \subseteq V$, rather than just connected ones), [14] shows that using $\tau = 0.5$ will maximize the expected classification accuracy over all vertices $v \in V$. In simulated instances of the connected ASD estimation problem (i.e. the Altered Subnetwork problem), we indeed see that using $\tau = 0.5$ yields a larger classification accuracy than the $\tau = \tau_{\text{NetMix}}$ (Figure S2). However, at the same time, for moderately small $\mu$ we see that using $\tau = 0.5$ also yields both a lower F-measure and a more inaccurate estimate of $|A|/n$, the fraction of nodes in the altered set (Figure S2).

#### 2.2 Using log-likelihoods in NetMix

In the NetMix algorithm, we make heavy use of the *responsibilities* $r_v = P(v \in A \mid X_v)$, which are used in the EM algorithm to fit a GMM to data. Another quantity which is often studied in the Gaussian mixture literature is the *log-likelihood ratio* $l_v = \log\left(\frac{P(v \in A \mid X_v)}{P(v \notin A \mid X_v)}\right)$. Note that $l_v > 0 \Leftrightarrow P(v \in A \mid X_v) > P(v \notin A \mid X_v)$.

In the NetMix algorithm, we use the estimated responsibilities to compute vertex weights as $w(v) = \hat{r}_v - \tau$. Another reasonable option is to instead use the estimated log-likelihood ratios in the weights, i.e. $w'(v) = \hat{l}_v - \tau$. Empirically, using responsibilities yields a slightly better performance compared to log-likelihoods (Figure S3). Moreover, one can easily generalize the responsibilities to account for multiple altered subnetworks (e.g. by using a GMM with more than 2 components), which is harder to do with log-likelihood ratios.

#### 2.3 Comparison between NetMix and heinz

The heinz algorithm [7] solves the following problem, which models vertex weights using a beta-uniform mixture (BUM) and requires a manual user-defined FDR parameter.

**Altered Subnetwork Problem (heinz [7] version).** *Let $G = (V, E)$ be a graph with vertex weights $\mathbf{X} = (X_v)_{v \in V}$ distributed as*

$$X_v \overset{i.i.d.}{\sim} \begin{cases} \text{Beta}(a, 1), & \textit{if } v \in A, \\ U(0, 1), & \textit{if } v \in V \backslash A, \end{cases} \tag{29}$$

*for some unknown $a \in (0, 1)$ and unknown* connected *subgraph $A \subseteq V$ of $G$. Given $G$, $\mathbf{X}$, and $\tau_{FDR} \in \mathbb{R}$, find a connected subgraph $C$ that maximizes $\sum_{v \in C} -(\log X_v - \tau_{FDR})$.*

We note that one can define the Altered Subset Distribution and its variants (unstructured, connected, etc.) using a beta-uniform mixture as above, and we refer to such a distribution as the BUM ASD.

There are two important distinctions between the normally distributed ASP and the problem that heinz is solving. First, heinz does not directly estimate the altered subgraph $A$, but rather identifies high-scoring connected subgraphs according to a value $\tau_{\text{FDR}}$ derived from a user-defined FDR. In the main text, we discuss how this manually tuned parameter can encourage users to selectively tune the FDR in order to influence which genes are included in the resulting altered subnetwork, i.e. "$p$-hacking". Indeed, published analyses using heinz use a large range of values of $\tau_{\text{FDR}}$. For example, [15] applied heinz to single cell RNA sequencing data and considers FDR thresholds as small as $10^{-26}$. [10] applied heinz to microarray data in

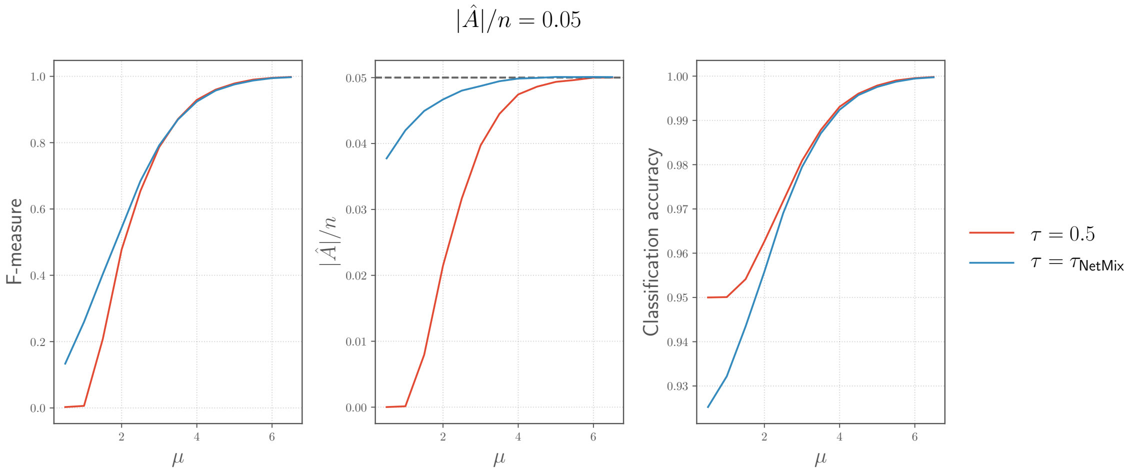


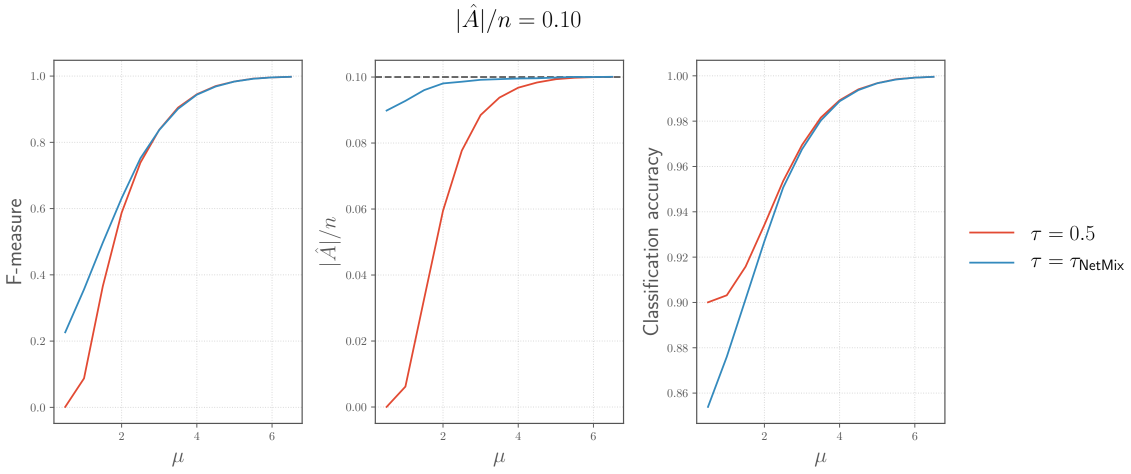


Figure S2: Comparison of NetMix with different choices of $\tau$ on simulated instances of the Altered Subnetwork Problem using the Hint+HI interaction network. The blue line represents the NetMix algorithm used in the main text.

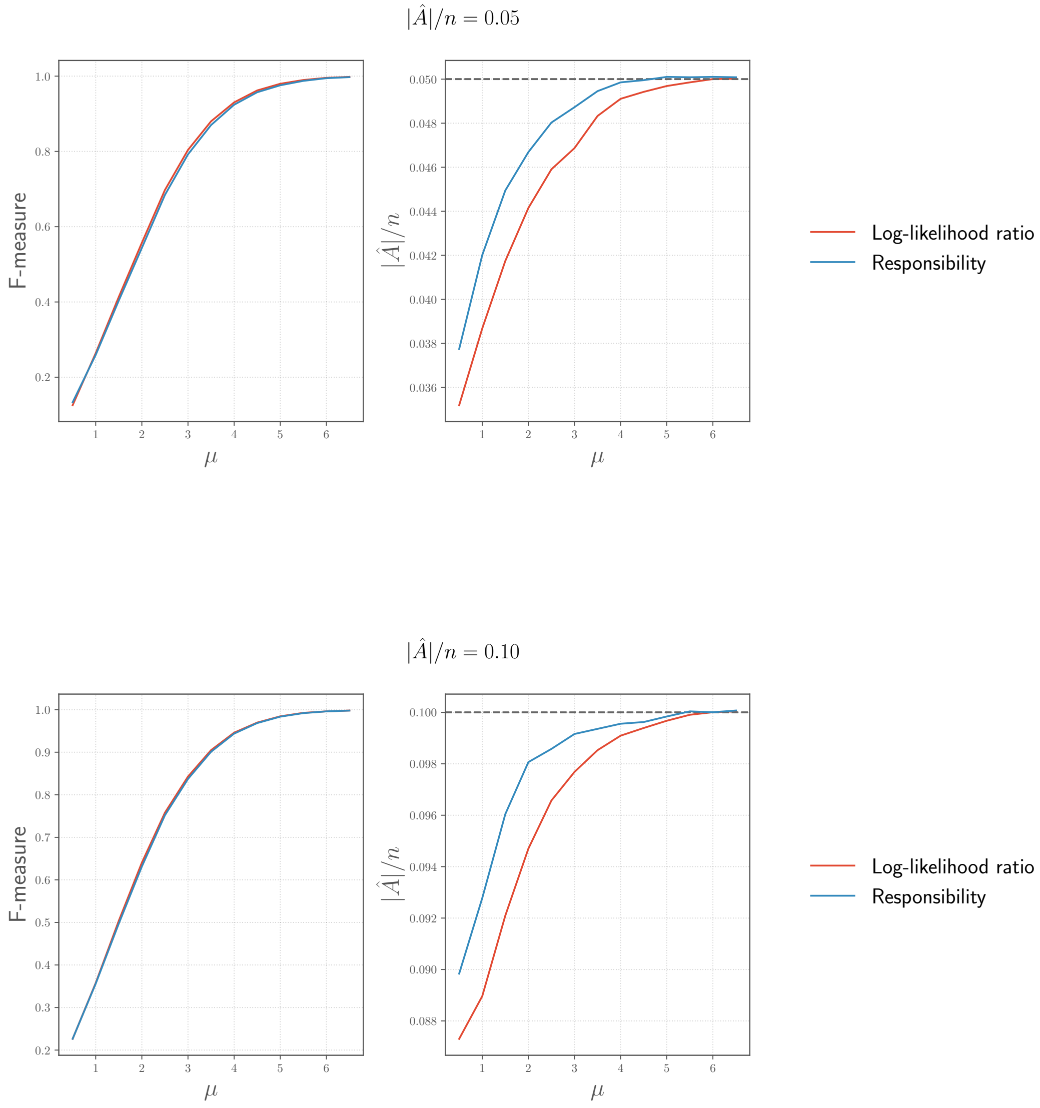


Figure S3: Comparison of NetMix using responsibilities versus log-likelihoods on simulated instances of the Altered Subnetwork Problem using the Hint+HI interaction network. The blue line represents the NetMix algorithm used in the main text.

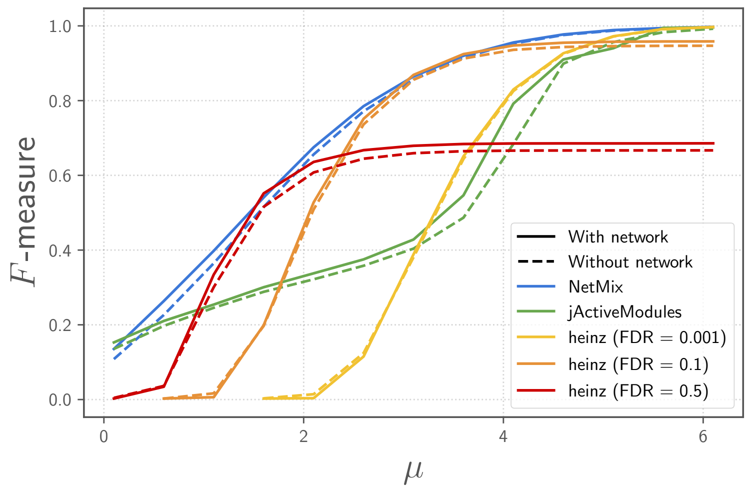


Figure S4: Comparison of altered subnetwork identification methods on simulated instances of the Altered Subnetwork Problem with and without the HINT+HI interaction network. Compare to Figure 4.

prostate cancer and "scanned a large range of FDRs to guarantee desirably sized modules and evaluated the obtained solutions in terms of recall and precision". [4] applied heinz to non-synonymous *de novo* mutations in early brain development and used a family-wise error rate threshold (FWER) threshold of $5 \cdot 10^{-6}$ derived from a Bonferroni correction, skipping the BUM model fit altogether. As we demonstrate in the results, determining the appropriate FDR value is a highly non-trivial task. We also demonstrate that, for nearly all choices of the FDR parameter, the heinz algorithm outputs a biased estimate of the anomaly size $|A|$, similar to jActiveModules (Figure S5).

Another distinction is that heinz models $p$-values with a beta-uniform mixture (BUM). The rationale for modeling $p$-values with a BUM model is purely empirical, with [18] stating that this mixture provides an "excellent approximation to the observed distribution of $p$-value" arising from microarray experiments in practice". [7] also writes that "modeling the signal component by a beta distribution is justified by [...] a Quantile-Quantile (Q-Q) plot". However, in a follow-up paper to [18], Pounds and Cheng [17] experimentally observe that the BUM is not a useful model in some settings, and that the BUM tends to underestimate the number of $p$-values drawn from the altered distribution. Interestingly, the simulations of [17] use normal distributions to generate their $p$-values.

### 3 Methods

#### 3.1 Data

##### 3.1.1 Gene scores

For our analysis of somatic mutations in cancer, we used MutSig2CV driver $p$-values [16] from the TCGA PanCanAtlas project [1] from https://gdc.cancer.gov/about-data/publications/pancan-driver as of October 1, 2019. We also used $p$-values for differential gene expression yeast from [13, 12]. Figure S6 shows the distribution of these gene scores, where we computed $z$-scores using the standard formula $z = \Phi^{-1}(p)$, where $p$ is a $p$-value and $\Phi$ is the CDF for the standard normal distribution.

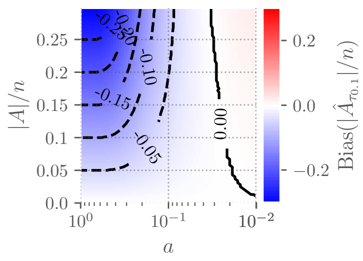

(a) Bias($|\hat{A}|/n$) for the unstructured BUM ASD using $\tau_{\text{FDR}}$ corresponding to FDR $= 0.1$.

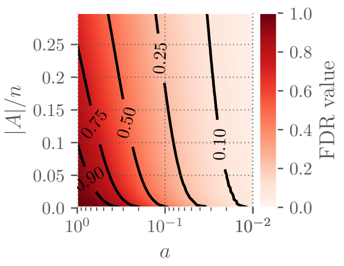

(b) FDR threshold needed to obtain an unbiased estimate of $|A|/n$ for the unstructured BUM ASD.

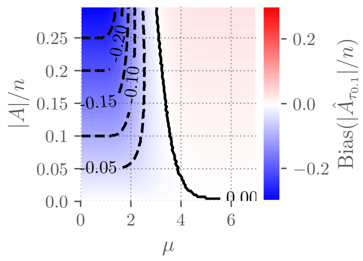

(c) Bias($|\hat{A}|/n$) for the unstructured (normally distributed) ASD using $\tau_{\text{FDR}}$ corresponding to FDR $= 0.1$.

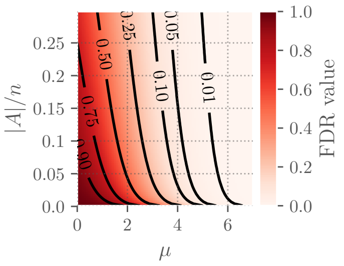

(d) Bias in estimate of $|A|/n$ for the unstructured (normally distributed) ASD using $\tau_{\text{FDR}}$ corresponding to FDR $= 0.1$.

Figure S5: Bias($|\hat{A}|/n$) for a fixed FDR, and the FDR threshold needed to obtain an unbiased estimate of $|A|/n$, for both the normally distributed and BUM.

##### 3.1.2 Interaction networks

For our analysis, we use the following interaction networks, which were the most recent versions available as of October 1, 2018.

- HINT+HI [6, 20]:
  - HINT binary
    `http://hint.yulab.org/download/HomoSapiens/binary/hq/`
  - HINT co-complex
    `http://hint.yulab.org/download/HomoSapiens/cocomp/hq/`
  - HuRI HI:
    `http://interactome.baderlab.org/download`
- iRefIndex 15.0 [19]:
  `http://irefindex.org/download/irefindex/data/archive/release_15.0/psi_mitab/MITAB2.6/9606.mitab.22012018.txt.zip`
- ReactomeFI 2016 [5, 8]:
  `http://reactomews.oicr.on.ca:8080/caBigR3WebApp2016/FIsInGene_022717_with_annotations.txt.zip`

For the ReactomeFI interaction network, we considered the set of interactions with a confidence score of 0.75 (out of 1) or larger. For each network, we treated each interaction as undirected, and we restricted our analysis to the largest connected component of the network.

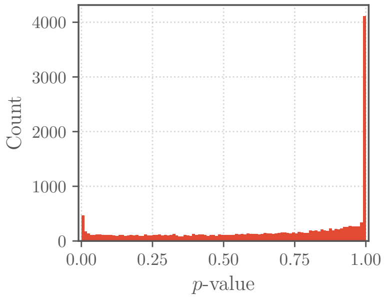


(a) MutSig2CV $p$-value distribution.

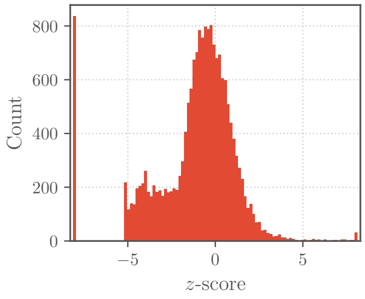


(b) MutSig2CV $z$-score distribution.

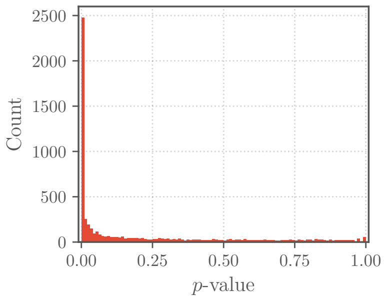


(c) *GAL1* $p$-value distribution.

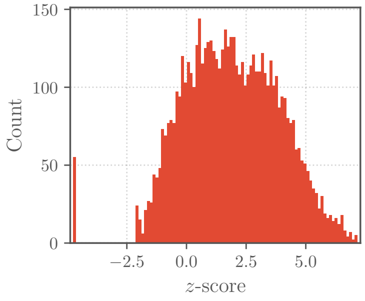


(d) *GAL1* $z$-score distribution.

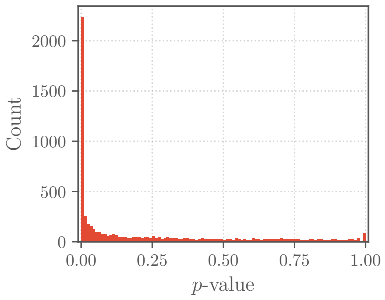


(e) *GAL4* $p$-value distribution.

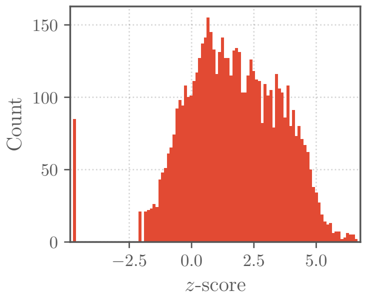


(f) *GAL4* $z$-score distribution.

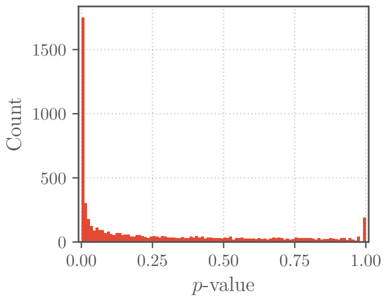


(g) *GAL80* $p$-value distribution.

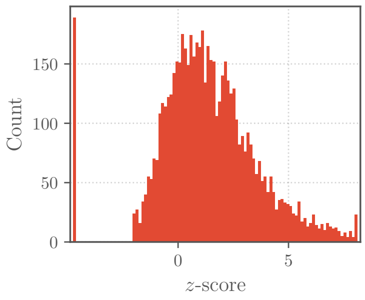


(h) *GAL80* $z$-score distribution.

Figure S6: Distribution of $p$-values for MutSig2CV driver $p$-values on TCGA PanCanAtlas somatic mutation data and differential gene expression $p$-values for GAL pathway alterations in yeast.

| Interaction network | Vertices | Edges | Density | M.D. | A.S.P. | Diameter |
|---|---|---|---|---|---|---|
| HINT+HI | 15,074 | 182,088 | $1.6 \cdot 10^{-3}$ | 11 | 3.4 | 9 |
| iRefIndex | 17,136 | 408,688 | $2.8 \cdot 10^{-3}$ | 21 | 3.0 | 8 |
| ReactomeFI | 11,501 | 209,020 | $3.2 \cdot 10^{-3}$ | 13 | 3.4 | 13 |

Table S1: Summary of the human interaction networks used in the analysis, where M.D. is median degree and A.S.P. is average shortest path.

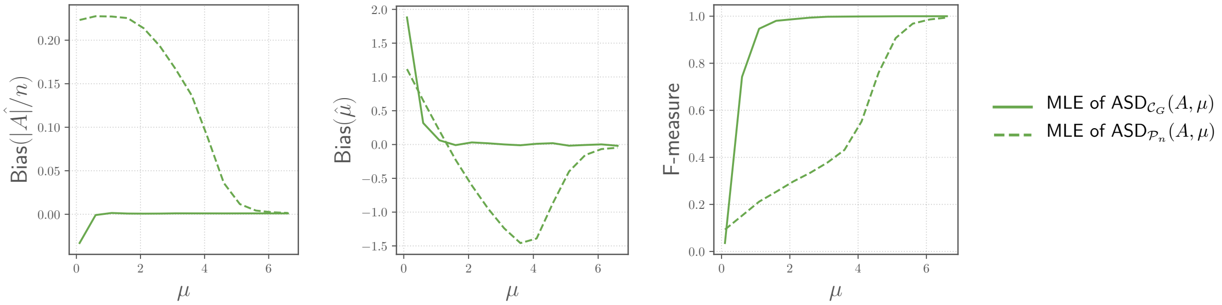


Figure S7: Comparing the MLE of the connected ASD $\text{ASD}_{\mathcal{C}_G}(A, \mu)$ for the connected line graph $G$ with $n = 1000$ vertices, to the MLE of the unconstraine ASD $\text{ASD}_{\mathcal{P}_n}(A, \mu)$, with altered subset size $|A| = 0.05n$.

#### 3.2 Running methods without a network

All three of the methods we run (heinz, jActiveModules*, and NetMix) find a subnetwork

$$\hat{A} = \underset{\text{connected } S \subset V}{\text{argmax}} \; f(S) \tag{30}$$

for some scoring function $f(S)$, e.g. jActiveModules* uses $f(S) = \frac{1}{\sqrt{|S|}} \sum_{v \in S} X_v$. For heinz and NetMix, the scoring function can additionally be written as $f(S) = \sum_{v \in S} w(v)$ for vertex weights $w(v)$. To run these methods without a network, we change each method to find $\hat{A}_{\text{no network}} = \text{argmax}_{S \subset V} f(S)$.

For jActiveModules, $\hat{A}_{\text{no network}}$ can be found efficiently by sorting the vertex scores $X_v$ and finding

$$\underset{k \in \{1,\dots,n\}}{\text{argmax}} \frac{1}{k} \sum_{i=1}^{k} X_{[i]} \tag{31}$$

where $X_{[1]} \geqslant X_{[2]} \geqslant \cdots \geqslant X_{[n]}$. For heinz and NetMix, we select the vertices $v$ with $w(v) > 0$.

#### 3.3 Comparison of other graph topologies

When using a biological interaction network, jActiveModules*, which computes the MLE of the ASD, has relatively poor performance in solving the Altered Subnetwork Problem (Figure 4). We hypothesized that this is because connectivity is a relatively weak constraint for a biological interaction network (i.e. most subsets of vertices are close to a connected graph). We test this by looking at another graph: the *line graph* $G$, where connectivity is a strong constraint (there are only $O(n^2)$ connected subgraphs of a line graph). For all $\mu$, the MLEs $\hat{A}_{\text{ASD}}, \hat{\mu}_{\text{ASD}}$ for the parameters of the connected ASD of the line graph, $\text{ASD}_{\mathcal{C}_G}(A, \mu)$, are much less biased (Figure S7).

| | Network | | | |
|---|---|---|---|---|
| Method | None | HINT+HI | iRefIndex | ReactomeFI |
| jActiveModules* | 2,136 / 0.155 | 1,575 / 0.191 | 1,815 / 0.174 | 557 / 0.261 |
| jActiveModules (Greedy search) | N.A. / N.A. | N.A. / N.A. | N.A. / N.A. | N.A. / N.A. |
| jActiveModules (Simulated annealing) | N.A. / N.A. | 12,284 / 0.086 | 15,046 / 0.074 | 8,329 / 0.118 |
| heinz (FDR $= 0.001$) | 115 / 0.205 | 119 / 0.216 | 109 / 0.217 | 114 / 0.215 |
| heinz (FDR $= 0.1$) | 259 / 0.244 | 249 / 0.264 | 259 / 0.255 | 253 / 0.215 |
| Hierarchical Hotnet | N.A. / N.A. | 228 / 0.214 | 297 / 0.215 | 228 / 0.214 |
| NetMix | 307 / **0.254** | 263 / **0.277** | 296 / **0.270** | 264 / **0.270** |

Table S2: Results of network methods on cancer driver gene prediction using MutSig2CV driver $p$-values from the TCGA PanCanAtlas project and multiple interaction networks. Each entry reports the size / $F$-measure of the altered subnetwork identified by each method.

##### 3.4 Somatic mutations in cancer

We ran several network methods on somatic mutation data in cancer using MutSig2CV $p$-values on TCGA PanCanAtlas somatic mutation data and the HINT+HI, iRefIndex, and ReactomeFI interaction networks.

We tried the greedy search version of jActiveModules, but it did not complete in under 100 hours on any network. We ran the simulated annealing version of jActiveModules cwith with default settings except with the number of iterations increased from 2,500 to 100,000. We tried fitting the MutSig2CV $p$-values using heinz's beta-uniform mixture model software `fitBumModel`, but it returned an error of `BUM model could not be fitted to data` on MutSig2CV $p$-values, so we used the Benjamini-Hochberg multiple testing correction [2] on the MutSig2CV $p$-values to set the parameter $\tau_{\text{FDR}}$ for FDR thresholds of 0.001 [7] and 0.1 [16, 11]. We ran Hierarchical HotNet, setting the heat scores of genes with MutSig2CV $q = 1$ to zero. We ran NetMix with the default settings, except genes with extremely significant gene scores (e.g., 30 genes have MutSig2CV $p < 2.2 \cdot 10^{-16}$) skew the fit of the altered and background distributions. Thus, we removed 115 genes with FDR $< 0.001$ before computing $\hat{\mu}$ and $|\hat{A}|/n$ but computed responsibilities for each gene, even genes with FDR $< 0.001$.

Table S2 summarizes the output of each network methods. Fig. S8 compates the outputs of different methods.

##### 3.5 Results on differential gene expression data from Expression Atlas

We compared the null proportions $\hat{\pi}$ estimated by both both the GMM (used by NetMix) and BUM (used by heinz), defined as follows:

$$\text{GMM} : \hat{\pi} = (1 - \hat{\alpha}),$$
$$\text{BUM} : \hat{\pi} = \hat{\lambda} + (1 - \hat{\lambda})\hat{a},$$

where $\lambda$ and $a$ are the mixing proportion and parameter to the Beta distribution respectively) [17].

In the main text, we note that many experiments from the Expression Atlas had $> 50\%$ of genes differentially expressed with FDR $\leqslant 0.1$. This tendency towards a large number of significantly differentially expressed genes is repeated in a number of other analysis. For example, Ideker et al. [12] analyzed *S. cerevisiae* gene expression following alterations in the galactose-utilization pathway using jActiveModules[13]. We found that *most* genes in these data were significantly differentially expressed using single-gene tests. In total, 3,212, 2,945, and 2,347 of the 5,934 measured genes had FDR $< 0.1$ [2] while 1,377, 1,088, and 834 genes had FDR $< 0.001$ for alterations in *GAL1*, *GAL4*, and *GAL80*, respectively. (see Figure S6 for distributions of these gene scores).

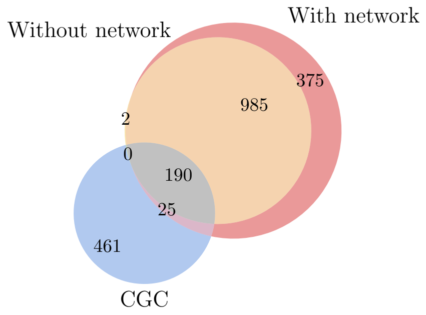


(a) jActiveModules* results

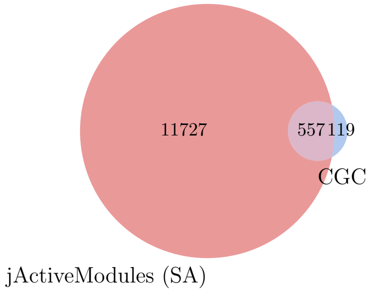


(b) jActiveModules results

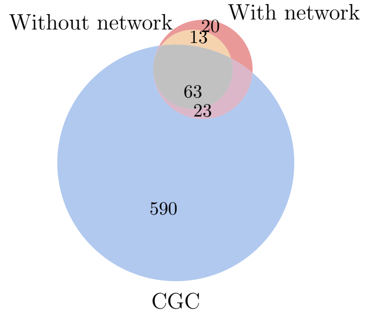


(c) heinz results with FDR = 0.001

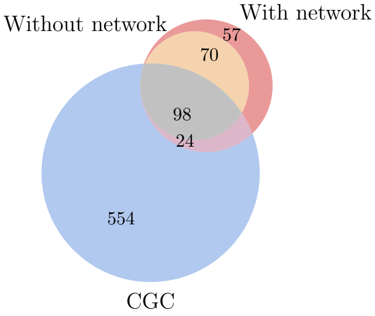


(d) heinz results with FDR = 0.1

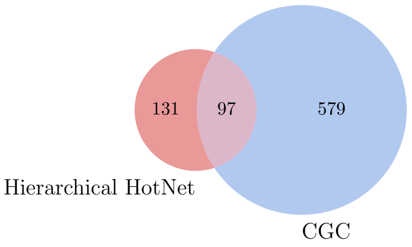


(e) Hierarchical HotNet results

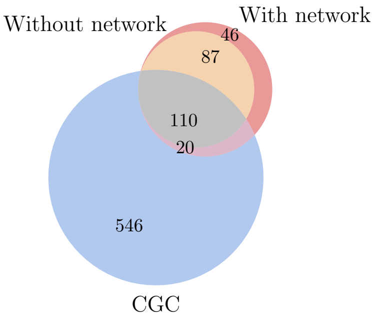


(f) NetMix results

Figure S8: Selected comparisons of results on MutSig2CV $p$-values for somatic mutations from the TCGA PanCanAtlas project on HINT+HI interaction network

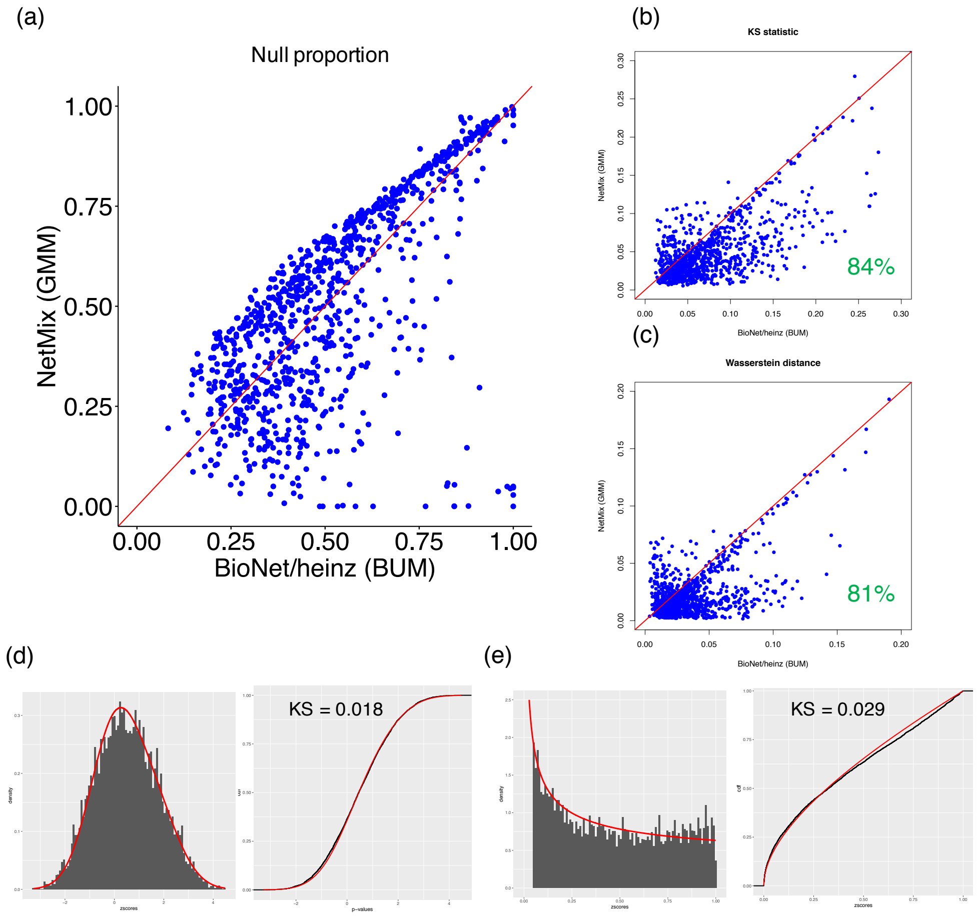


Figure S9: (a) Null proportion of 945 experiments from the Expression Atlas fit by GMM (y-axis) and BUM (x-axis) (b) Wasserstein distance between empirical p-values and GMM fit (y-axis) vs. BUM fit (x-axis) (c) KS statistic between empirical p-values and GMM fit (y-axis) vs. BUM fit (x-axis) (d) GMM fit to transformed p-values from E-MTAB-2973 (e) BUM fit to p-values from E-MTAB-2973.

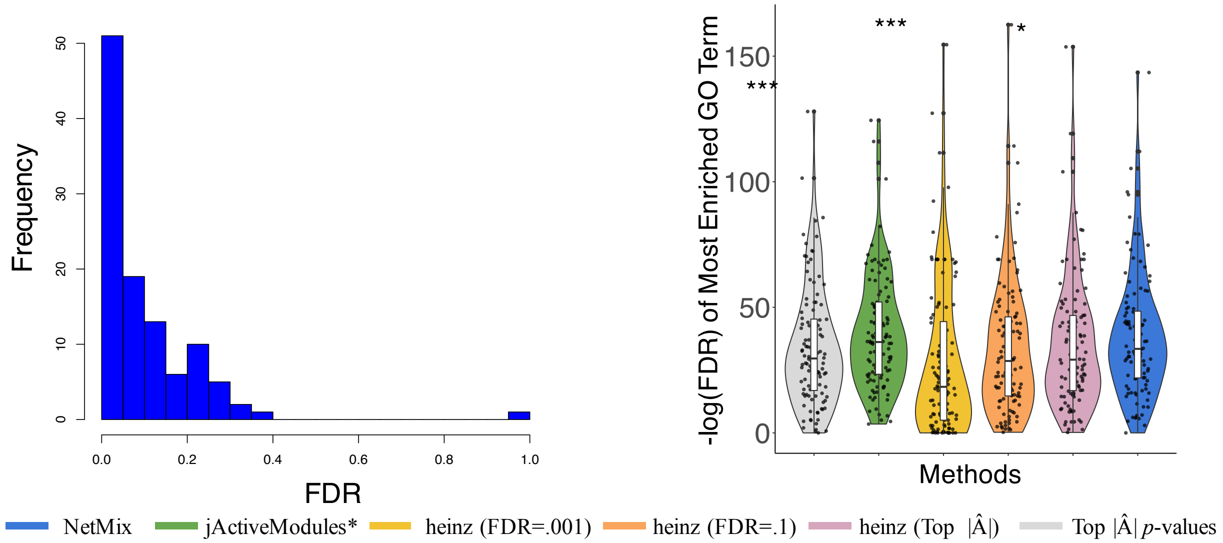


Figure S10: (left) Histogram of FDR thresholds used for heinz/BioNet such that the number of postive scored nodes was matched the size $|\hat{A}|$ of the alternative subset predicted by NetMix. (right) $-\log(p$-value) for the most enriched GO term for gene subsets estimated by each method using the HINT+HI network.

[9] Jason Gaitonde. Big-o bounds on the k-th largest element of iid gaussians (answer), cs theory stack exchange. `https://cstheory.stackexchange.com/questions/44161/big-o-bounds-on-the-k-th-largest-element-of-iid-gaussians`.

[10] Hao He, Dongdong Lin, Jigang Zhang, Yu-ping Wang, and Hong-wen Deng. Comparison of statistical methods for subnetwork detection in the integration of gene expression and protein interaction network. *BMC Bioinformatics*, 18(1):149, 2017.

[11] Heiko Horn, Michael S Lawrence, Candace R Chouinard, Yashaswi Shrestha, Jessica Xin Hu, Elizabeth Worstell, Emily Shea, Nina Ilic, Eejung Kim, Atanas Kamburov, et al. Netsig: network-based discovery from cancer genomes. *Nature methods*, 2017.

[12] Trey Ideker, Owen Ozier, Benno Schwikowski, and Andrew F Siegel. Discovering regulatory and signalling circuits in molecular interaction networks. *Bioinformatics*, 18(suppl 1):S233–S240, 2002.

[13] Trey Ideker, Vesteinn Thorsson, Jeffrey A Ranish, Rowan Christmas, Jeremy Buhler, Jimmy K Eng, Roger Bumgarner, David R Goodlett, Ruedi Aebersold, and Leroy Hood. Integrated genomic and proteomic analyses of a systematically perturbed metabolic network. *Science*, 292(5518):929–934, 2001.

[14] Adam Tauman Kalai, Ankur Moitra, and Gregory Valiant. Efficiently learning mixtures of two gaussians. In *Proceedings of the Forty-second ACM Symposium on Theory of Computing*, STOC '10, pages 553–562, New York, NY, USA, 2010. ACM.

[15] Florian Klimm, Enrique M Toledo, Thomas Monfeuga, Fang Zhang, Charlotte M Deane, and Gesine Reinert. Functional module detection through integration of single-cell rna sequencing data with protein–protein interaction networks. *bioRxiv*, page 698647, 2019.
